# Age distributions of rare lineages reveal recent demographic history and selection

**DOI:** 10.1101/265900

**Authors:** Alexander Platt, Jody Hey

**Affiliations:** Temple University, Center for Computational Genetics and Genomics, Philadelphia, PA 19122

## Abstract

The age of an allele of a given frequency offers insight into both its function and origin, and the distribution of ages of alleles in a population conveys significant information about its history. The rarer the allele the more likely it is to reveal functional biological insight and the more recent the historical revelation. By measuring the length of the haplotype shared between an individual carrying a rare variant and its closest relative not carrying the variant we are able to approximate the age of the variant and can apply this method even when only a single copy of a variant has been sampled in a population. Applying this technique to rare variants in a large population sample from the United Kingdom, we identify historical migration from West Africa approximately 400 years ago, evidence of direct selection against novel protein-altering rare variants in individual biological pathways, continued negative frequency dependent selection on protein-altering variants in olfactory transducers and the innate immune system, and map the impact of background selection on the most recent portions of the sample genealogy.

## Introduction

The age of an allele can reveal much about how it came to its observed frequency. In particular an allele that is younger than expected given its frequency is likely to have been under directional selection. This effect is not surprising for evolutionarily favored alleles, but it also applies to the case of harmful alleles^1^. Such alleles are often quickly lost from a population, but those that persist and turn up in a sample of genomes are likely to be young. In this context “harmful” refers to Darwinian fitness, but it is probable that many alleles with negative impacts on health would also be under negative selection^2,3^. Thus, if we estimate allele age, we can combine this with other information such as allele frequency, geographical distribution, and functional annotation to improve predictions of an allele’s effect on human health.

Alternatively, an allele that is older than expected given its frequency is also a candidate for having an interesting history. Alleles that are maintained in the population by balancing selection can be much older than their frequency alone would suggest^4,5^. Admixture can also be a source of alleles that are older than they appear to be on the basis of allele frequency alone^6^.

In developing an estimator of allele age, we considered several constraints. The estimator should not be a function of allele frequency, it should not be a function of the demographic history of the sampled population, and it should not be highly sensitive to the details of the data pipeline. We also wish to be able to study the age of the very rarest alleles, including those that appear only once in a sample (singletons). This last criterion rules out methods that are based on the similarity of different copies of an allele at flanking sequences^7,8^. Such methods can inform on the time at which different copies of an allele last shared a common ancestry, which is a useful proxy for allele age^8^, but they cannot be applied to singletons.

We turned instead to an approach that utilizes comparisons between a chromosome carrying a focal allele and other chromosomes not carrying that allele. Consider an alignment of two chromosomes, one with and one without a derived allele at a SNP, and consider that the genealogy or gene tree at the site of the allele will coalesce when the two chromosomes most recently shared a common ancestor for the base position of the SNP. Then a search that extends along the chromosome to the right or the left will reveal a distance to the first flanking mismatch between the chromosomes. This mismatch will have been caused by a mutation on one of the two gene tree edges, or by a variant introduced by a recombination event on one of those edges. This distance can be modeled as a Poisson process with a rate parameter that is the product of the time of common ancestry and the sum of recombination and mutation rates per base position per generation^9–11^. By extension if we compare the chromosome with the allele with all other chromosomes not carrying that allele, we can identify that which shows the longest flanking region without a mismatch, what we call the maximal shared haplotype or *msh*. This will be a function of first coalescence time, *t_c_*, between the chromosome with the allele and all chromosomes not carrying the allele.

We developed a coalescent model for the probability density of *t_c_* as a function of *msh*. For a singleton, the model provides a likelihood estimator, 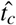, that uses both *msh* values (to the 5’ and 3’ directions from the focal base). For alleles that occur more than once, it can be used as a composite likelihood estimator of time of first coalescent event ancestral to the edge upon which the mutation causing that allele occurred. The bias for 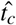 is low for rare alleles, including singletons, and it works similarly well when using sequenced chromosomes with known or estimated phase, despite the presence of switch errors in the latter (figure S1 and table S1). Having age estimates for singleton alleles also provides a means for estimating the phase of singletons when estimating chromosome haplotypes from short-read data (fig S3). Though the method assumes a demographic model for the length distribution of the sister to the edge carrying the mutation, the resulting estimator is quite insensitive to this model (figure reffig:robust demography).

For identifying harmful alleles, 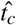 will become increasingly useful as sample sizes grow, as the degree to which rare alleles are enriched for alleles under directional selection will be higher the rarer an allele is. Our focal data set, the UK10K genome sequencing study of over 7,200 genomes (3,600 individuals), has singleton alleles at an estimated population frequency less than 0.00014. We first asked if the distribution of 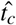 values is consistent with previously estimated demographic histories for the UK, and then turned to questions of the direct selective effects of rare alleles, and finally to a study of indirect selective effects (background selection).

The probability that any two haplotypes share a common ancestor in a particular generation depends on the size and structure of the population. In small populations there are fewer ancestors overall and therefore a greater opportunity to share one than in a larger population, and in subdivided populations two individuals from different subpopulations are less likely to share a common ancestor than individuals from the same subpopulation. These kinds of processes do not depend on the function (or lack thereof) of any particular genetic locus and are simply properties the movements of individuals. Therefore, they simultaneously influence the ages of coalescence events all across the genome. Any given demographic model of the history of a population implies a particular distribution of coalescence times for neutral loci everywhere in the genome. To assess the ability of recently published demographic histories of Europe and the United Kingdom we compare their predicted distributions of times of coalescence (*t_c_*) with those estimated at loci associated with rare variants in the UK10k population sample. We also use this distribution to fit additional parameters explicitly modeling potential recent immigration from Africa to the United Kingdom.

Natural selection, by contrast, acts to perturb the *t_c_* distribution heterogeneously across the genome. Natural selection can act directly on individual genetic variants to alter the expected length of the lineage on which they are found compared to neutral variants of the same frequencies. While neutral variants may persist at low frequencies for extended periods, or return to low frequencies after drifting to higher frequencies, deleterious rare variants are being steadily removed from the population and are prevented from drifting to higher frequencies or sojourning there. This leads to a younger age distribution of rare deleterious variants than neutral variants. Negative frequency dependent selection, where rare alleles confer an advantage over common ones, on the other hand, preserves rare variants (thus preventing their loss) and makes common variants rarer. Both of these phenomena will contribute to an older *t_c_* distribution associated with rare, negative frequency dependent variants than rare neutral variants. Previous work seeking to identify genetic variants undergoing current or recent selection have focused on relatively common variants (*e.g.*^12,13^). These can show signals of strong positive selection where new or very rare variants have quickly risen to high frequencies, and directional selection acting on common variants contributing to highly polygenic traits. Using *t_c_* estimates we can see the direct influence of natural selection as it influences rare variants.

## Results

### Neutral variation and demographic history

#### Age distribution of rare variation in United Kingdom

We calculated 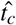 for 21,992,410 of the rarest variants in the UK10k^14^ whole-genome population sequencing sample that has been filtered to remove close relatives and individuals of non-European ancestry. The distribution of estimates revealed a dramatic excess of variation that is both old and rare – well beyond what is predicted by previous models of UK or European human history. Figure 1 shows the means and standard deviations of the distributions of log(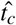) values for variants found 2, 3, 4, 5, 10, and 25 times in the UK10k sample of 3,621 individuals (7,242 haplotypes), and compares them with predictions from five published models of UK and European demographic histories^15–19^ as well as new models with additional admixture events from an African population or diverged archaic human group. For all of the lowest frequency classes, the observed data contain variants that are far too old to have been generated by the published models (all of which returned mean simulated *t_c_* distributions considerably smaller than for 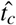 from the UK10K singletons). The models proposed by Gutenkunst et al.^15^ and Gravel et al.^16^, the two published models with migration between Africa and Europe, return standard deviations similar to that of the observed data but with insufficient old alleles to substantially raise the predicted mean. For variants as common as those found 25 times (a frequency of 0.35%), all models fit the observed distribution reasonably well.

**Figure 1.**
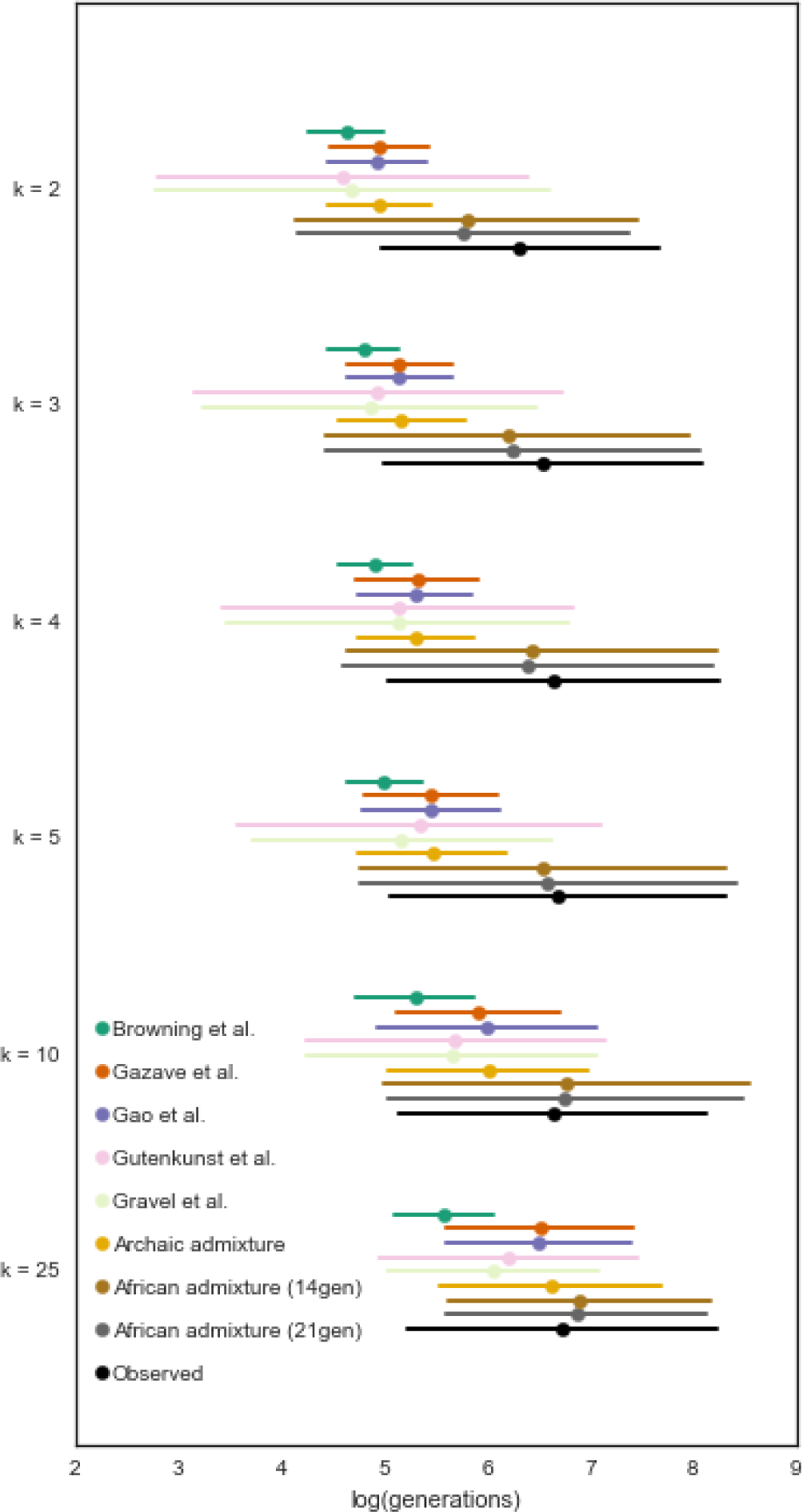
Distributions of log(*t_c_*) values. Variants of different frequencies in the UK10k data (*k* values) represented as mean ± one standard deviation of the log-transformed *t_c_* values. The observed distribution is marked in black with other colors indicating expectations under proposed demographic histories of Britain and Europe.

Admixture from archaic humans will have introduced old alleles, and some of these are expected to appear at the lowest frequencies in the UK10K sample. However, we found that admixture with archaic humans does not introduce sufficiently rare alleles in the numbers necessary to explain the discrepancy. While the alleles introduced by such admixture are old, few of the introduced alleles end up in the relevant frequency classes at the time of sampling. When we turned to extant human populations as potential sources of old, very low frequency variants, we found an excellent fit with models that include recent admixture from African populations.

Figure 2 shows the fit of a series of models based on that by Gazave et al.^17^ with the addition of a recent migration event from an un-sampled African population to an ancestral UK population from which it separated 2,000 generations ago. The best fit model is one with 1.2% admixture 21 generations ago. This is a model in which 10% of rare UK10k variants predate the human expansion out of Africa (see extended data figure S5); have been segregating at moderate frequency within Africa; and were recently introduced to the UK population from Africa through migration.

**Figure 2.**
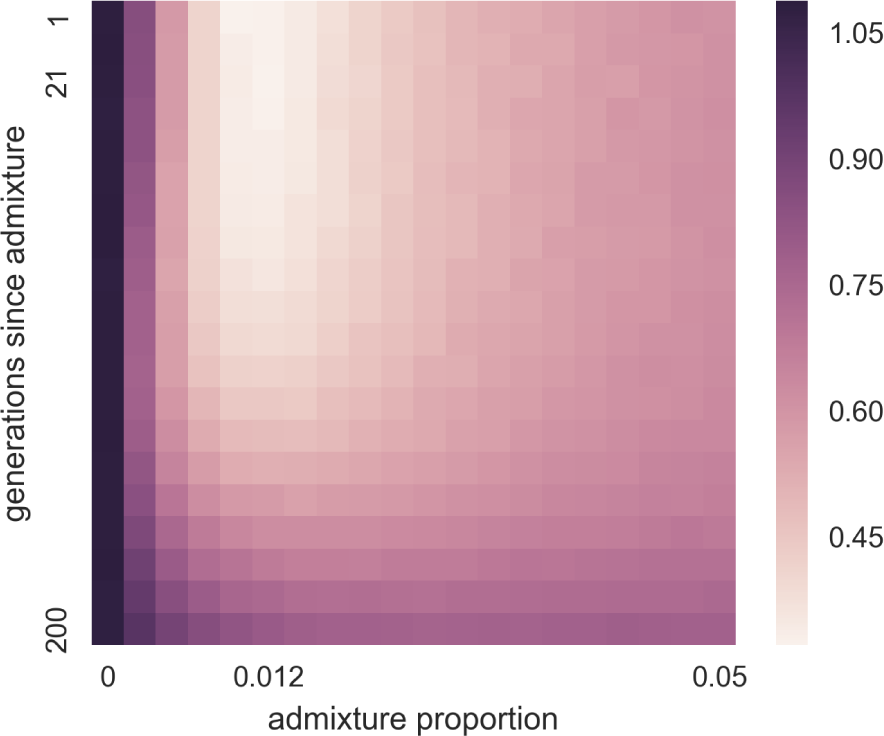
African admixture parameter optimization. Root mean squared log error between observed *t_c_* values and those simulated over a grid of proposed values for timing and magnitude of African gene flow into the ancestors of the UK10k population.

#### Distribution of old rare UK variation across geography

If migration from Africa is responsible for substantially altering the age distribution of rare UK alleles we expect to find a substantial proportion of these rare British alleles at higher frequency in African populations than they are in the UK (or other European populations). We assessed all of the UK10k singleton variants, and variants found 25 times, for their presence and frequency within the populations of the 1000 genomes project phase 3 dataset^20^. Figure 3 shows that very rare UK10K alleles are typically at their rarest in Great Britain and present in higher frequencies in African populations than elsewhere in the world. In contrast, variants observed 25 times within the UK10k data show a very different pattern, as they are rarely found outside of Europe, but are found at relatively high frequency when they do occur. This is the pattern expected of alleles that have primarily not been recently introduced by migration, but rather have persisted both inside and outside of Africa since their origins before the population divergence at the time of the out of Africa expansion.

**Figure 3.**
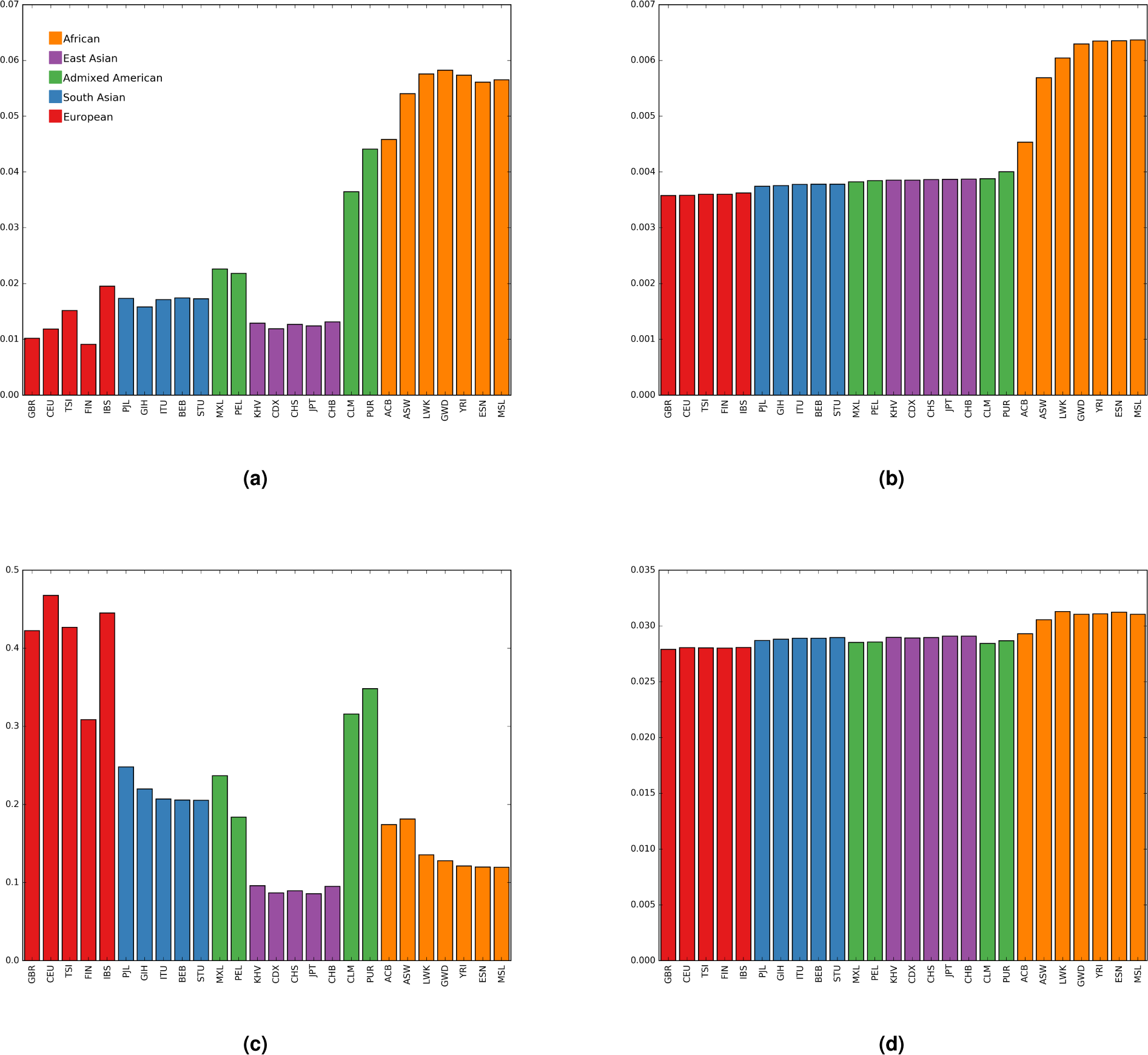
Geographic distribution of UK10k variants. Panels (a) and (b) reflect variants that are singletons in the UK10k data. Panels (c) and (d) reflect variants found 25 times in the UK10k data. Panels (a) and (c) describe the proportion of UK10k variants found in each population sample in 1000 Genomes Project data. Panels (b) and (d) describe the average frequency in the 1000 Genomes Project data of UK10k variants. 95% confidence intervals are all less than 0.4% of the bar heights for panels (a) and (b) and less than 4% of the bar heights for panels (c) and (d)

Extended data figure S6 shows that the distributions of 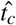 values for UK10k variants that are found in African populations in the 1000 Genomes Project data are considerably older than those that are not.

#### Distribution of old rare variation across UK individuals

While figure 2 indicates support by the data for models with 1.2% admixture, the overall distribution of rare allele ages is consistent with a range of models that vary in the timing of the admixture event. We further refined the estimate of the time of admixture by explicitly modeling the distribution of quantity of introgressed alleles observed among the individuals in the UK10K sample. In a random-mating model with admixture occurring over a short period of time, the introgressed alleles will come to be spread fairly evenly in the population over a small number of generations. Figure 4 shows that the distribution of the number of doubleton alleles with 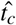 ages predating the out of Africa expansion shows considerable clustering across individuals, with 13% of individuals harboring 40% of all such variants. This clumping is suggestive of recent introgression, but could also be consistent with older introgression where the decay of clustering slowed by non-random mating. As shown in figure 2, the observed distribution is well fit by a broad range of models of assortative mating, all implying a time of admixture 11-14 generations before sampling. The model ‘African admixture (14gen)’ in figure 1 represents the expected distribution of log(*t_c_*) values for a 1.2% admixture event 14 generations before sampling.

**Figure 4.**
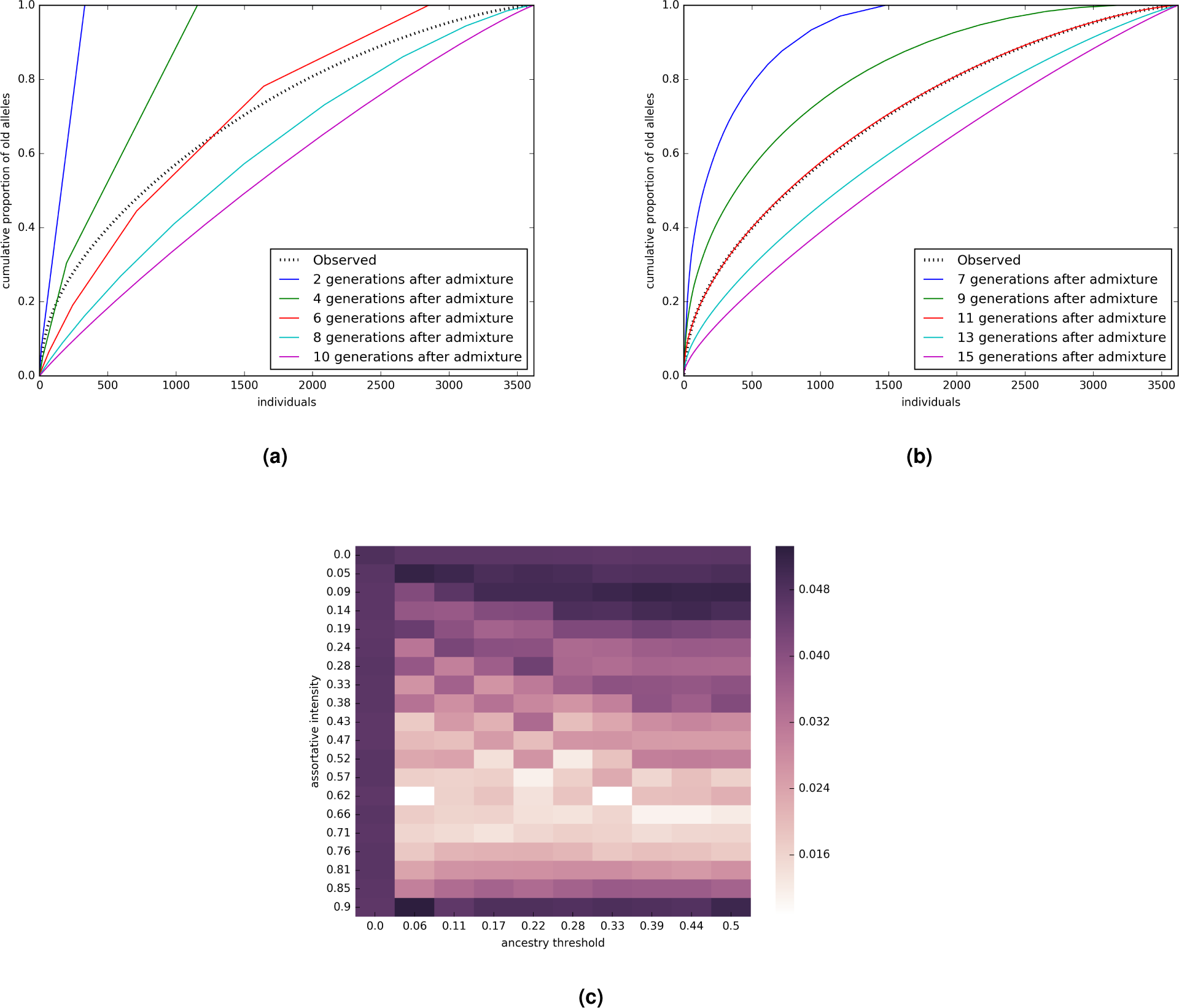
Cumulative proportion of doubleton alleles more than 2,500 generations old carried by individuals ordered from most to fewest old doubleton variants. Observed distribution is marked in dotted black. Colored lines are simulations of 1.2% admixture at different historic time points and assuming (a) random mating or (b) assortative mating with a threshold of 22% and intensity of 63%. Panel (c) shows the minimum (across ages) root mean square error between the observed cumulative proportion of old doubletons and that predicted under a range of models of non-random mating. Dark bars at the top and left indicate poor fit of random mating models. Light bar across the middle indicates similar support for models with a wide range of ancestry thresholds.

If Africa populations are indeed the source of many of the old singletons in the UK, then individuals carrying large numbers of old singletons should show higher sharing of non-singleton alleles with African populations. We tested this prediction by assessing the differential similarity between UK10K genomes (that have either high or low counts for singletons with estimated ages older than 3,000 generations) with an ABBA-BABA test^21,22^. For each of UK10K genomes the *D* statistic was determined using a random CEPH genome (representative of Europe) and a random Yoruban genome (representative of Africa) from the 1000 genomes dataset^20^. The two distributions of *D* statistics show some overlap (extended data figure S7). However the UK10K genomes with low rare singleton counts each had mean *D* values (overall mean of-0.3615) lower than each of the high rare singleton count genomes (overall mean of −0.3500) consistent with closer proximity to Yoruban genomes for the high singleton count UK10K genomes. A *t* test of the two groups of mean *D* values was highly significant (*t* = 5.9589, d.f. = 8, *p*=0.0003).

### Direct selection on rare variants

With the expectation that protein-altering genetic variants will experience greater purifying selection than other genetic variants, we contrasted the distribution of *t_c_* values for UK10k singleton missense, nonsense, and splice-site variants with singleton variants that are not protein-altering. As shown in figure 5, the mean log(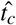) value for the 1e5 protein-altering singleton variants is 5.72 log(generations), with the mean log(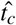) value for the other 1.5e7 singleton variants being 5.85 log(generations), (Wilcoxon-Mann-Whitney *p <* 1e-128).

**Figure 5.**
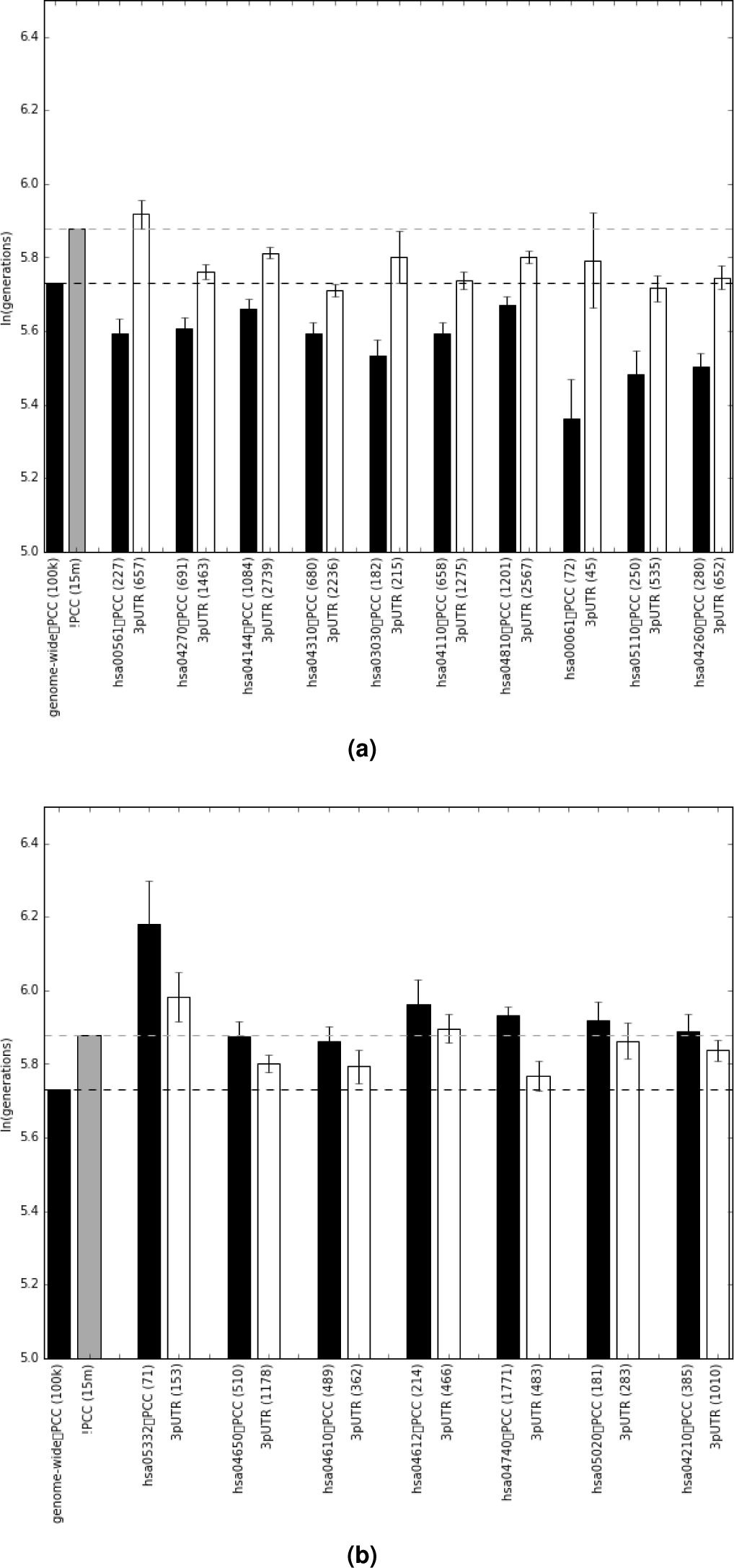
Mean log(*t_c_*) values for singleton variants for each KEGG pathway with non-zero *β_PC×KEGG_* coefficients: (a) are negative, (b) are positive. Dashed lines indicate mean log(*t_c_*) values for all protein-changing (black) and non-protein changing (grey) variants.

Natural selection is expected to act heterogeneously across genes contributing to different phenotypes. In highly constrained genes new variation will be deleterious and quickly removed leaving an increased signature of younger protein-altering singletons than singletons in the 3-prime untranslated region (3’UTR). Genes experiencing overdominance or other form of negative frequency-dependent selection will exhibit the opposite signature with singleton protein-altering variants being older than corresponding variants in the 3’UTR. We used KEGG^23^ pathway annotations for each gene as cofactors in a single LASSO^24^ regression to identify pathways independently contributing to gene-specific variation in the difference between protein-altering singleton variations and those in the 3’UTR. This resulted in 16 pathways with non-zero coefficients. Nine have positive coefficients causing protein-altering variants to be younger than expected given their 3’UTR variants: hsa00561 (Glycerolipid metabolism), hsa04270 (Vascular smooth muscle contraction), hsa04144 (Endocytosis), hsa04310 (Wnt signaling pathway), hsa03030 (DNA replication), hsa04110 (Cell cycle), hsa04810 (Regulation of actin cytoskeleton), hsa00061(Fatty acid biosynthesis), hsa05110 (Vibrio cholerae infection), and hsa04260 (Cardiac muscle contraction). Seven have negative coefficients making protein-altering variants older than expected given they ages of the 3’UTR variants: hsa05332 (Graft-versus-host disease), hsa04650 (Natural killer cell mediated cytotoxicity), hsa04610 (Complement and coagulation cascades), hsa04612 (Antigen processing and presentation), hsa04740 (Olfactory transduction), hsa05020 (Prion diseases), and hsa04210 (Apoptosis). Figure 5 shows the mean log(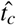) values for the protein-altering and 3’UTR singletons in each of these pathways.

### Indirect selection on external lineages

As a general estimator 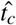 is versatile in some ways that might not initially be appreciated. In particular the estimator is easily modified to estimate the time of first coalescence for any external edge of the genealogy at any base position. By simply not including the probability of a mutation in the *msh* likelihood, we can adapt the estimator and obtain edge lengths for each individual chromosome for any base position in the genome. We applied the estimator in this way, and measured the geometric mean edge length at even megabase intervals and within each gene for a total of 20,261 invariant points in the aligned genomes of the UK10K genomes, measurements we refer to as 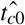. As these measurements are not taken at positions supporting polymorphisms, and because all positions are similarly subject to the shared demographic history of the UK population sample, the primary source of variation in 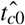 will be the indirect selective effects of linked alleles. In other words, we can use 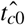 as a new tool for the study of background selection.

We find a striking pattern of variation in 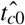 along chromosomes. If background selection had the effect of rescaling external branches of gene trees the same way it rescales the total length of gene trees (as is assumed when modeling background selection as a reduction in effective population size^25^, 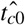 should be directly proportional to *B*, a measure of local gene tree rescaling due to background selection inferred from population polymorphism levels^26^. Normalizing the 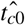 values to have the same mean as *B* values yields a statistic *α* that is directly comparable to *B* (figure 6). We find that *α* is more variable overall than *B* (variance of 0.074 vs. 0.057), is only moderately correlated with *B* (Pearson’s *r* = 0.27), and exhibits higher levels of autocorrelation than *B* over distances less than four megabases but less autocorrelation at distances greater than four megabases.

**Figure 6.**
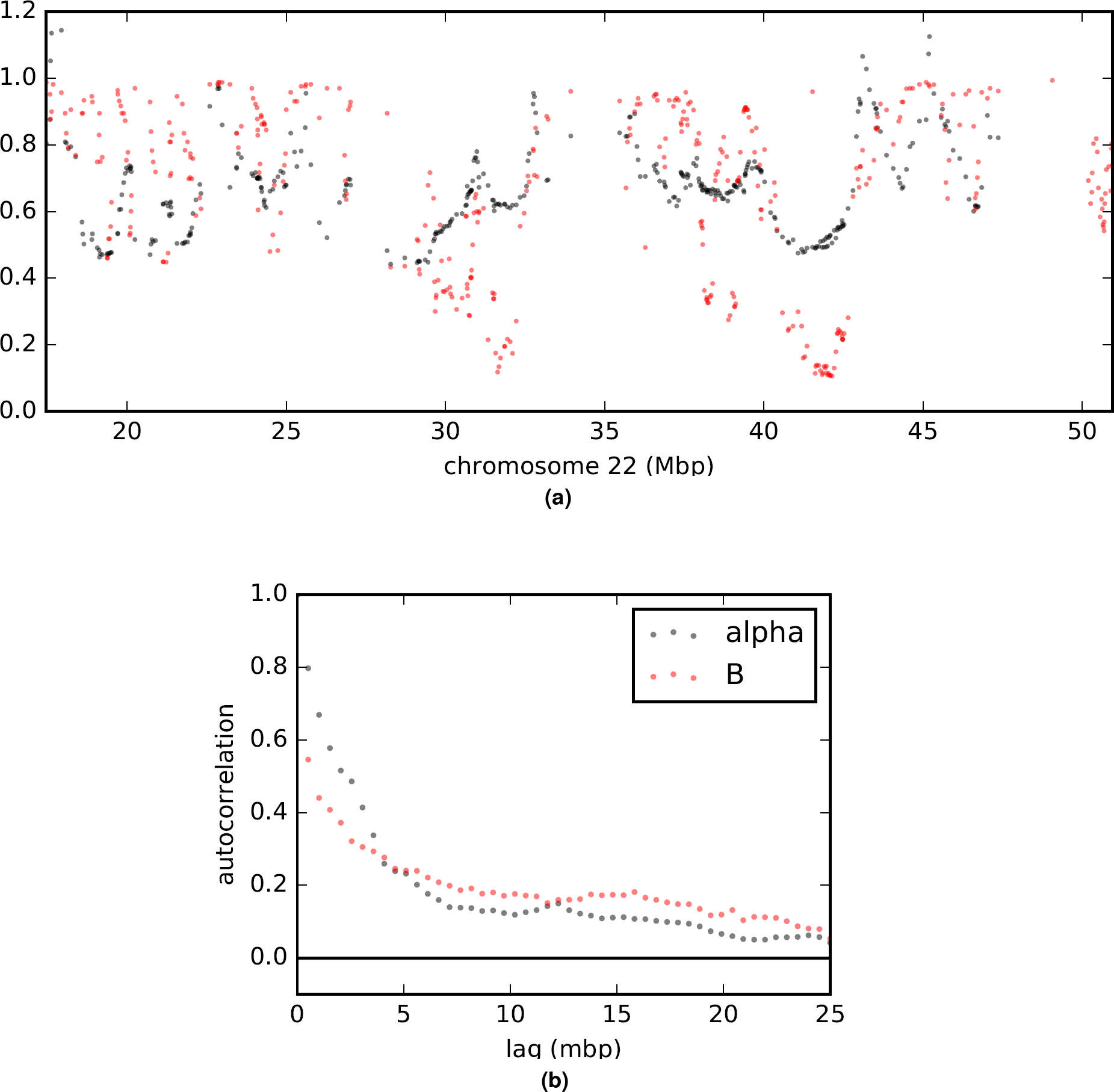
Comparison of *a* (rescaled 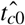; black) and *B* (in red) values across the genome. (a) is a plot of measurements arrayed along a representative chromosome. Other chromosomes are shown in extended data figures S8t169-S14. (b) is an autocorrelation plot. With between 20,258 and 17,610 paired observations per point, all values shown are significantly greater than zero by Pearson’s moment correlation test with a maximum *p <* 1.5e-8.

## Discussion

Coalescent models connecting patterns of genetic variation to statements of genealogy structure and depth have long provided the statistical underpinning for much of population genomics^27^. He were develop a coalescent-based estimator of genealogy edge length that can be used to address important questions on selection and admixture while also being robust to demographic history and to important sources of error in genome assembly and phasing. For derived alleles carried by one or multiple chromosomes, 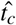 can be used to identify alleles that are unusually young or old, relative to others of the same frequency, and it can be used to estimate age distributions for classes of alleles. The estimator can also be used to study the length of the external edges for the genealogy of a large sample at any point in the genome, thereby providing information on the recent history of indirect selective effects on gene tree depths.

With African gene flow modeled as a point admixture event, we estimate 1.2% of the ancestors of the UK10k population came from Africa 11-14 generations ago. A generation time of 29 years^28^ places the migration event early in the era of the “First” British Empire when Britain was actively establishing colonies in West Africa and the West Indies^29^. During this period there was a notable rise in the number of African individuals in the UK Black British population, often as household attendants to returning sea captains and colonists or as former slaves from Spain and Portugal^30,31^.

This model of demographic influx is one in which many alleles that arose before the expansion out of Africa rose to (or remained at) relatively high frequencies within a much larger African population while being lost or excluded from the smaller ancestral European population. These alleles then had the opportunity to be reintroduced to the European population in very small quantities through recent migration. We are not excluding the possibility that the gene flow that shifted the distribution of UK10k *t_c_* values was a more complicated phenomenon than a single migration event. With relatively higher levels of African gene flow into Southern European populations^32,33^ it is probable that some proportion of the African variants we find in the UK10k population did not come by immigration directly from Africa but by more circuitous means.

It has been argued that rare variants should fall almost exclusively into the class of recent mutations, and that rare variants are unlikely to be found outside of their population of origin^16,34^. This conclusion was primarily drawn from a model fit from a smaller sample where the rarest variants were at considerably higher population frequencies than the rarest variants in the UK10k data. Studies explicitly designed with individuals of admixed ancestry have also noted elevated quantities of rare variants^6^. As shown in figure 3, even in a sample screened to reduce non-European contributions, as a variant rises in frequency even to 0.0035 frequency (*k*=25 in the UK10k sample), the relative probability of finding it outside of Europe drops dramatically. It is predominantly among the very rarest of variants that we see this substantial impact of recent migrants.

Human genetic variation is being shaped both by ongoing selective constraint as well as ongoing balancing/negative frequency dependent selection, with large parts of the innate immune system experiencing the latter. The effects of selective constraint can be seen throughout the genome. The geometric mean of 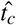 for protein-changing singletons is 50 generations younger than singletons that do not change proteins. This 16% difference is strongly conservative given the rate of haplotype phasing errors that homogenize these two groups.

Above and beyond this genome-wide selective constraint removing protein-changing mutations from the population, individual biological pathways show unique signatures of selection acting on rare protein-changing variants. The pathways showing especially elevated selective pressures reducing *t_c_* values of rare protein-changing variants are extremely diverse, ranging from metabolic pathways to cell signaling pathways to DNA replication. Together they comprise a broad set of genes for which any alteration is most likely to have health and fitness consequences.

Pathways with protein-changing singleton 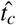 values that are older than expected (older, even, than for non-protein changing singletons) fall into two categories: olfactory transduction and components of the innate immune system. The unique structure of olfactory receptor cells, wherein each cell expresses only a single olfactory transduction gene, has led to an hypothesis that heterozygosity in olfactory transduction genes may contribute to a larger palette of sensory responses^35^. If such an expanded palette contributed to evolutionary fitness it would create an overdominance effect that would be expected to retain rare functional variants. Our results suggest that selection to retain rare functional olfactory alleles^36,37^ is not a phenomenon relegated to the deep evolution of the human species but something that has persisted into our recent history. That multiple different pathways related to the innate immune system all show a non-zero LASSO coefficient tells us that even though the pathways overlap in their comprising genes, the negative frequency dependent selection causing the signal is not isolated within a small number of core genes. Rather, broad swaths of this entire system are under selective pressure to retain rare functional variants.

Analysis of 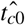 values demonstrates a discordance between the effects background selection has had on deeper ancestral portions of gene trees and recent, external branches. Total population diversity reflects the size of the total gene tree, and estimates such as *B* ^26^ of the effects of background selection based on population diversity are strongly sensitive to variation in the lengths of long ancestral branches. The signal we see in the variation in 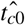 (or *α*) is only moderately correlated with *B* and has a different spatial structure. That *α* exhibits a different autocorrelation function than *B* indicates that background selection not only rescales external branches by a different amount than internal branches, but the entire scope of what linked sites are contributing to background selection, and the degree to which sites of a particular strength of linkage contribute, varies between internal and external branches.

## Methods

### UK10k data preparation

We use the mapping sample of 3,621 individuals from the UK10k data set that has been filtered by the UK10k consortium to remove close relatives and individuals of recent non-European ancestry. These include genomes from the ALSPAC cohort, which focused on the Avon region, and the TWINSUK cohort which includes samples from across the UK. We masked all CpG / TpG transversion polymorphisms to avoid homoplasy and reduce heterogeneity in the mutation rate. Haplotype phase for variants found two or more times was inferred by the UK10k consortium using SHAPEIT^38^.

Haplotype phase for singleton variants was assigned to maximize their *t_c_* values as branches of gene trees that harbor mutations are expected to be longer than those that do not. As shown in extended data figure S3, coalescent simulation confirms that this is a highly effective method for phasing singleton variants when the difference between the *t_c_* values for the two possible phases is large.

### Estimating *t_c_*

We developed an estimator of the coalescent time of a chromosome, *t_c_*, at a particular base position that is a function of the maximum shared haplotype (*msh*) lengths. Starting at that base position and comparing the focal chromosome to all other chromosomes, we identify the longest perfect match before the first discrepancy. This is done independently in the 5’ and 3’ directions giving a pair of *msh* observations that need not arise from the same alignment.

Under the assumptions of the Sequential Markov Coalescent (SMC), the length of shared haplotype extending from a point (whether it carries a SNP or not), between any pair of chromosomes, can be modeled as a function of their coalescent time and the sum of mutation rate and recombination rate^9^. For the case of the *msh* identified in comparisons with all chromosomes, we show by simulation (extended data figure S4) that the closest mutation or recombination event is well approximated as having occurred on either the external branch leading to the singleton variant (which create discrepancies between the variant-carrying haplotype and all other haplotypes in the sample) or on its first sister branch (which create discrepancies between the variant-carrying haplotype and all of its closest relatives). Each *msh* is then treated as the distance to the closest event in a Poisson process with a density parameter equal to the sum of the mutation and recombination rates times the sum of two branch lengths: the external branch along which the singleton variant arose, which has length *t_c_*, and its sister branch, which has some length *φ*. With a uniform per-base recombination rate *ρ* and mutation rate *µ*, the joint density of 5’ and 3’ *msh* values is the product of two exponential distributions, each of the form

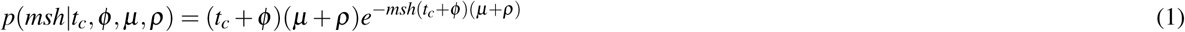

Apart from *ϕ*, all of these terms are either known (*msh*,*μ*,*ρ*) or are to be estimated (*t_c_*). In order to integrate over *ϕ* we adapt a hazard model of the rate of coalescence forward in time, to develop a probability density for *ϕ* conditional on *t_c_*. The number of branches ancestral to a sample that existed *t* generations before the sample was taken can be approximated as

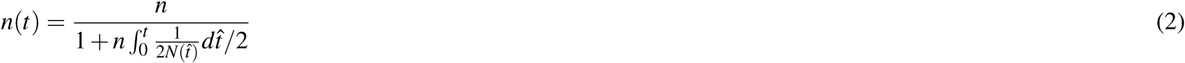

where *n* is the sample size and *N*(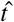) is the historical effective size of the population 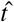 generations before sampling^7^. By substituting *t* = *t_c_* − *ϕ* in equation 2, we reverse the time direction, and by taking the derivative with respect to *ϕ* we have an expression for the instantaneous rate of coalescence in the gene tree as time moves forward:

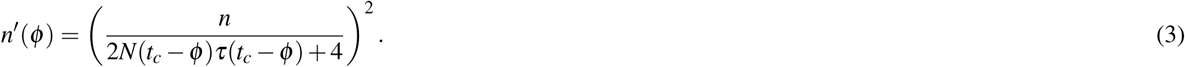

The instantaneous rate of coalescence along one particular branch, our hazard function *λ* (*ϕ*), is the rate for the tree divided by the number of branches:

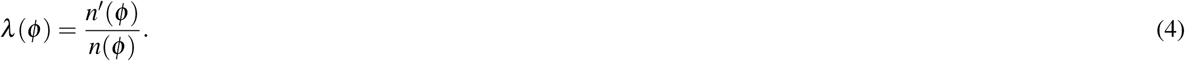

The probability that the next coalescent event to happen on the branch occurs after *ϕ* generations is the product of the probability of a coalescent event at generation *ϕ* and the probability of zero coalescent events in generations 0 through *ϕ*:

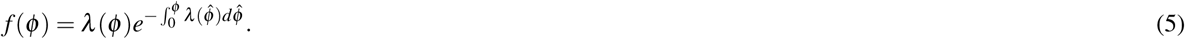

As a branch length, *ϕ* is bounded such that if no coalescent event happens within *t_c_* generations, the branch terminates at time zero instead of at a coalescent event. This leaves the conditional probability distribution as

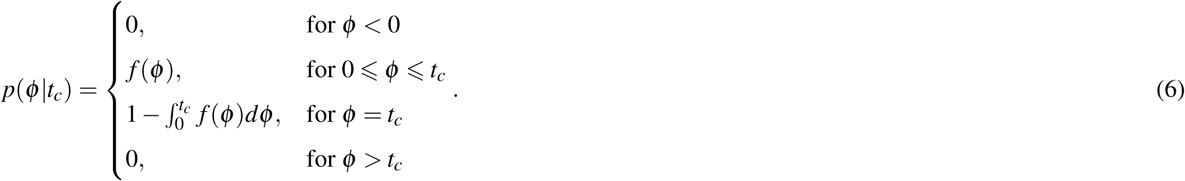

Then integration over *ϕ* yields the likelihood

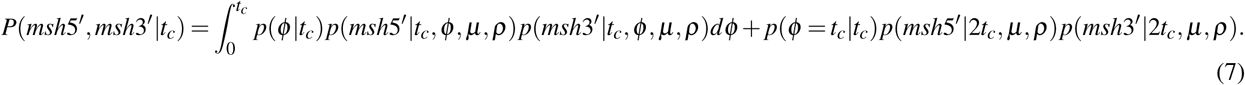

If the focal chromosome has a derived allele at the base position, then this likelihood also includes a term for the probability of at least one mutation in *t_c_* generations: (1 − *e*^−*μt_c_*^). The entire expression is readily simplified for a constant or exponential growth population model, while more complex demographic histories may require numeric integration. In either case the maximum likelihood estimate, 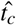 can readily be found by numerical optimization.

In practice we relax the assumption of uniform recombination rate. For a given SNP we record *χ* = *μ*(*x*_5′_ + *x*_3′_ + *c*_5′_ + *c*_3′_), where *x*_5′_ and *x*_3′_ are the *msh* distances in base pairs on the 5’ and 3’ sides of the focal base position and *c*_5′_ and *c*_3′_ are the same distances in Morgans. Then following^9^ we replace the product of exponential densities in (7) with

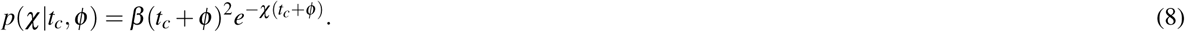

Here *χ* = (*μ*(*msh*5 = +*msh*3^′^) + *c*_5′_ + *c*_3′_) and *b* = (*μ* + *ρ*_5′_)(*μ* + *ρ* _3′_), where *c*_5′_ and *c*_3′_ are the distances in Morgans spanned by *msh*5^′^ and *msh*3^′^, respectively, and *ρ*_5′_ and *ρ*_3′_ are the recombination rates per base pair at the respective distal ends of the *msh* tracts.

With a detailed genetic map, and a set of aligned and phased genomes, *χ* values are easily measured. However our model assumes that *msh* will end precisely at the point of a mutation or recombination event, when in fact, in the case of recombination, such events cannot be directly observed, and so the measured *msh* necessarily continues until a (non-singleton) mutation is encountered that is not shared between the aligned haplotypes. This is expected to occur shortly after a recombination event as non-shared mutations accrue at a much higher rate beyond the first recombination event. We also wish to to minimize mistakenly shortened *msh* values due to mis-phased singleton variants, which can be quite common, so we do not end a maximum shared haplotype tract if a discrepancy is due to a singleton variant.

Because the demographic model enters (7) through the density of *ϕ*, which makes a minority contribution to the likelihood, the choice of the form of the demographic model (e.g. constant versus exponential versus multi-phase) actually has very little impact on parameter estimates (see extended data figure S2). This insensitivity of the estimator to demographic assumptions is a useful property of a statistic that is to be used to uncover components of demographic histories. We use a fixed *N* = 10,000 for all analyses of the UK10K variants.

For variants that are not singletons, where we wish to estimate the *t_c_* value shared by the different copies of an allele, we obtain an estimate by optimizing a composite likelihood function that is the product of the likelihoods of the *msh* values for each copy of the variant considered independently. In this case the *msh* values for each copy of an alleles are obtained by comparing that chromosome to all others that do not carry that allele.

While statistics derived from maximum values of observed data (such as the maximum length of a shared haplotype in our estimator) are often poorly behaved with large biases and strong sensitivities to model misspecification or stochastic noise, our estimator performs quite well, even under adverse conditions, such as when using *msh* values from statistically phased haplotypes. We demonstrate this through coalescent simulations generated with msprime^39^. As shown in Extended Data Table 1 and Extended Data Figure 4, for all of our simulations the bias was low. Precision was good for simulated populations of constant size but was low when estimating *t_c_* for very young singleton variants in models of extreme recent population growth. However even in these cases, the bias is small and precision may be sufficiently good to allow studies of many variants. For non-constant population size models precision returns to the higher levels found in constant-sized populations for variants of slightly higher frequencies.

It is possible that difficulties may arise if attempting to apply this estimator to very high frequency variants, however. If *n* (*t_c_*) is very small the approximation in equation 2 may behave poorly and require a different expression. That equation 1 need only consider *t_c_* and *ϕ* has only been confirmed for rare variants and modeling events on further parts of the tree may be necessary for common variants. Finally, extremely old variants may have *msh* values that approach the typical inter-polymorphism distance measured between unrelated individuals making accurate measurement difficult.

### Evaluating demographic model fit

We estimate the mean and standard deviation for a given model by simulating 500 independent coalescent trees with sample sizes of 7,242 haplotypes using msprime. For a low frequency allele observed *k* times we identify all of the branches that are ancestral to *k* leaves and calculate the average of their starting ages weighted by the ratio of their lengths to the total length of branches with *k* leaves across the set of trees.

We evaluated the distributions of *t_c_* values under eight different demographic scenarios. Browning et al.^19^ is a single-population model fit to UK10k data. Gazave et al.^17^ and Gao et al.^18^ are single-population models fit to the European-American population of the NHLBI Exome Sequencing Project^40^. Gutenkunst et al.^15^ and Gravel et al.^16^ are three-population models (African, European, and Asian) fit to data from the Environmental Genome Project^41^ and 1000 Genomes Project respectively, with all of the samples treated as coming from the European population. Archaic admixture is a model of ours that augments Gazave et al. with an archaic population that diverged 25,000 generations in the past and contributed 3% to the European population 2,000 generations ago. Our models of African admixture (African admixture (21gen) and African admixture (14gen)) use the Gazave et al. demographic model but add an additional 1.2% contribution of African lineages to the European population 21 or 14 generations ago respectively. Model fit is ascertained by calculating the root mean squared error of the means of the log(*t_c_*) values for variants occurring 2, 3, 4, 5, 10, and 25 times in the UK10K data.

### Estimating model parameters

Parameters describing the timing and quantity of African admixture were ascertained in a two-stage process. First, root mean squared log error of the mean *t_c_* values for variants occurring 2, 3, 4, 5, 10, and 25 times in the UK10K data were evaluated for each cell of a 20 x 20 two-dimensional grid of parameter values, with admixture varying between 0 and 5% and the time of admixture varying from 1 to 200 generations prior to sampling. Second, we refined our estimate of the time of admixture by modeling the spread of immigrant doubleton alleles through the population. At the time of admixture, all of the immigrant alleles will reside in a proportion of the population equal to the admixture proportion. In subsequent generations these alleles spread across more individuals. Using the admixture proportion of 1.2% identified in the previous grid search, we fit a time since admixture using a Wright-Fisher forward simulation with a two-parameter model of associative mating. A threshold parameter defines two ancestry groups and an intensity parameter describes the degree of preference for within-group mating. In each generation the population is divided into two ancestry groups: individuals with more than the threshold African ancestry and those with less. For each offspring in the next generation, one parent is drawn at random from the population. With probability equal to the intensity parameter the second parent is chosen at random from within the same ancestry group as the first parent. With probability one minus the intensity parameter the second parent is chosen at random from the entire population. Each offspring is assigned an ancestry proportion equal to the mean of its parents’.

### D stats

We calculated the *D* statistic (ABBA-BABA test^21^) for individuals carry high or low numbers of old singleton alleles (defined as alleles with estimated ages over 3000 generations) with respect to Yoruban genomes (representative of Africa) and CEPH genomes (representative of Europe) from the 1000 genomes project^20^. UK10K individuals were ranked by the number of singletons with estimated *t_c_* time over 3,000 generations. The highest 5 individuals and the lowest 5 individuals were each compared in ABBA-BABA tests with a Yoruban genome and a CEPH genome. A mean *D* was determined for each of the 10 UK10K genomes in 64 tests using all pairs of CEPH and Yoruban genomes (8 each) randomly sampled from the 1000 genomes data set.

### Annotation analysis

Every variant was assigned a protein changing status from its SnpEff^42^ annotation. Variants annotated as missense, stop gained, stop lost, start lost, splice acceptor, or splice donor, were categorized as protein changing. All others were considered non-protein changing.

To assess the effects different biological pathways have on the age of singleton variants we fit a regression analysis to each singleton variant that is either protein changing or found in the 3’ untranslated region (3’UTR) of a gene. Every variant *i* has an associated binary indicator *x_PCi_* that is 1 if it is a protein changing variant and 0 if it is a 3’UTR variant, and a vector of binary indicators 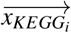 identifying all of the KEGG pathways to which the gene in which it is found belongs. We use the model

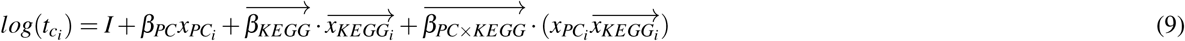

where *I* is an intercept, *β_PC_* is the amount by which protein changing variants differ from non-protein changing variants, 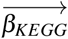 is the amount by which all variants (protein changing and 3’UTR) locally deviate within each pathway, and 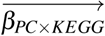 is the amount by which protein changing variants have pathway-specific deviations with respect to comparable 3’UTR variants. We fit all *β* parameters using the least absolute shrinkage and selection operator (LASSO)^24^ which minimizes the function

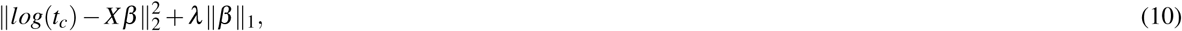

where *λ* is a regularization coefficient optimized through ten-fold cross validation. The regularization shrinks coefficients to zero whenever possible. This prevents overfitting while allowing the simultaneous analysis of many correlated factors. Where two factors are highly correlated the one with the strongest predictive power is retained and the other set to zero. Whenever two correlated factors are both assigned non-zero coefficients it is due to independent effects of each factor meaningfully contributing to the distribution of *t_c_* values.

### Background selection analysis

Estimates for *B* ^26^, the relative total length of a local gene tree compared to its hypothetical length if not subjected to background selection^43,44^, were downloaded from http://www.phrap.org and lifted over to hg19 coordinates. 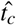 estimates were generated for every chromosome at monomorphic loci distributed at megabase intervals as well as at the start of every gene. Loci that did not have a reported *B* value were omitted. The 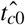 value at each locus is the geometric mean of the values from each chromosome. These 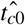 values were multiplied by a constant to produce *α* values with the same mean as the *B* values. Autocorrelation was calculated as the paired Pearson’s *r* coefficients of each locus and the next locus at least the indicated number of base pairs away.

## Acknowledgements

Funding was provided by NIH Grant RO1GM078204 to J.H.

## Author contributions statement

Both authors contributed to all stages of this research and manuscript.

---

**Table S1.** Performance characteristics of *t_c_* estimator. Properties of estimated *t_c_* values compared to true simulated *t_c_* values. a) attributes of variants found *k* times in the sample. b) simulations of either constant-sized populations or one with recent exponential growth. c) haplotype phasing taken directly from the simulation (known) or statistically inferred from simulated diploid genotypes. d) root mean squared log error of estimator. e) average signed log error. f) Pearson’s *r* statistic.

| $k^a$ | demography<sup>b</sup> | phasing<sup>c</sup> | error<sup>d</sup> | bias<sup>e</sup> | correlation<sup>f</sup> |
| --- | --- | --- | --- | --- | --- |
| 1 | constant | known | 0.16 | 0.09 | 0.87 |
| 5 | constant | known | 0.11 | 0.04 | 0.84 |
| 10 | constant | known | 0.10 | 0.02 | 0.82 |
| 25 | constant | known | 0.11 | -0.02 | 0.69 |
| 1 | constant | inferred | 0.39 | -0.16 | 0.41 |
| 5 | constant | inferred | 0.11 | 0.03 | 0.82 |
| 10 | constant | inferred | 0.10 | 0.01 | 0.81 |
| 25 | constant | inferred | 0.11 | -0.03 | 0.68 |
| 1 | growth | known | 0.17 | 0.11 | 0.55 |
| 5 | growth | known | 0.10 | 0.06 | 0.76 |
| 10 | growth | known | 0.09 | 0.04 | 0.85 |
| 25 | growth | known | 0.09 | 0.01 | 0.81 |
| 1 | growth | inferred | 0.22 | -0.05 | 0.26 |
| 5 | growth | inferred | 0.10 | 0.03 | 0.70 |
| 10 | growth | inferred | 0.09 | 0.02 | 0.82 |
| 25 | growth | inferred | 0.09 | -0.01 | 0.80 |

**Figure S1.**
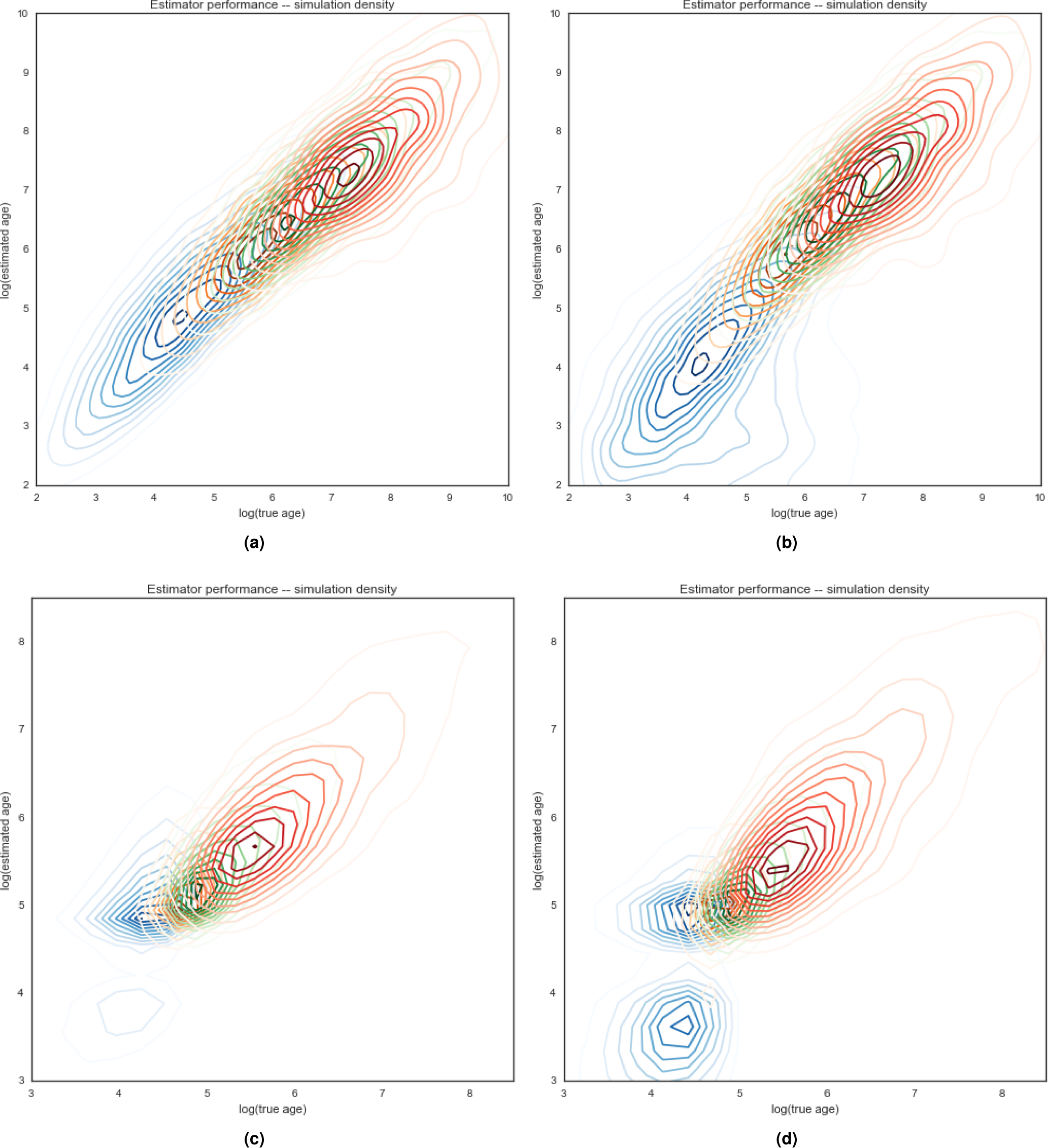
Estimator performance. Density plots for estimation of *t_c_* values as a function of true *t_c_* values in simulated data. Panels (a) and (b) are simulated constant-sized populations of 10,000. Panels (c) and (d) are populations that have grown exponentially from 10,000 to 500,000 over the last 120 generations. Panels (a) and (c) are evaluated on true known haplotypes. Panels (b) and (d) use statistically phased haplotypes from simulated genotypes. Densities of singleton variants is indicated in blue, variants sampled five times in orange, 10 in green, and 25 in red.

**Figure S2.**
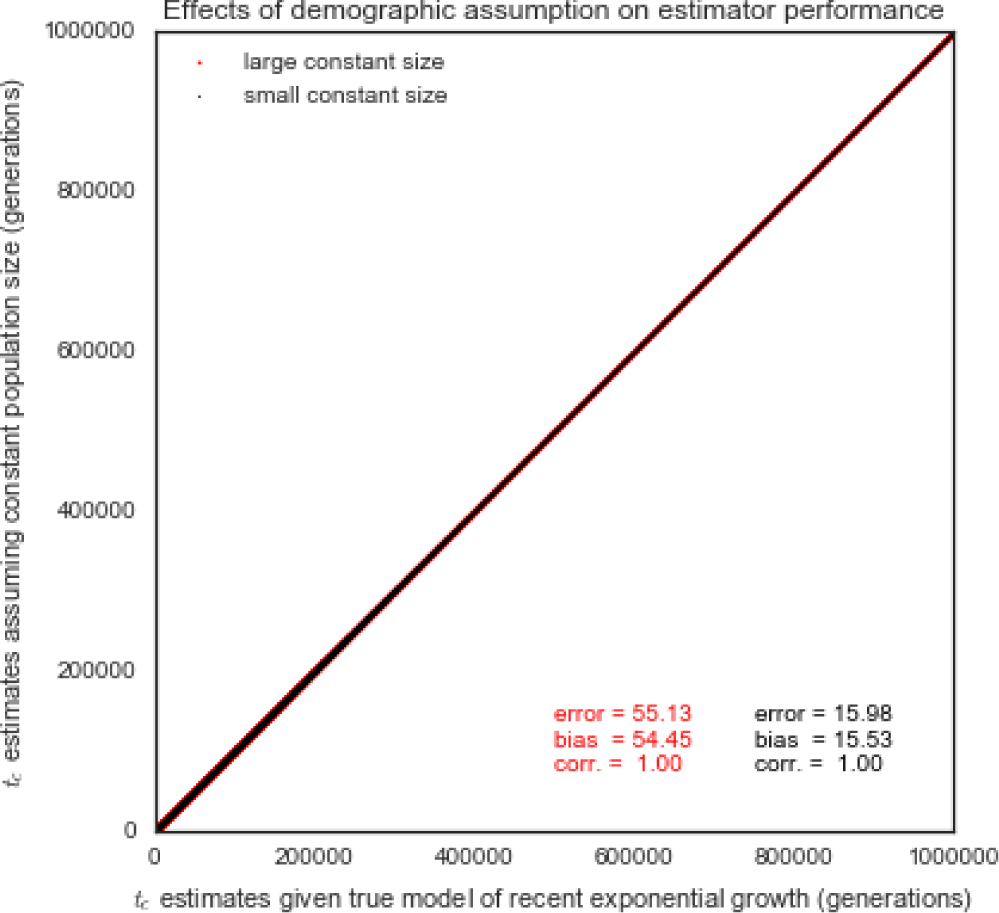
Estimator robustness to demographic model assumptions. Distribution of estimated *t_c_* values for simulated singleton variants estimated assuming a constant-sized population of 10,000 (black) and 500,000 (red) as a function of the value estimated assuming the correct generative model of exponential population growth from 10,000 to 500,000 over the last 120 generations.

**Figure S3.**
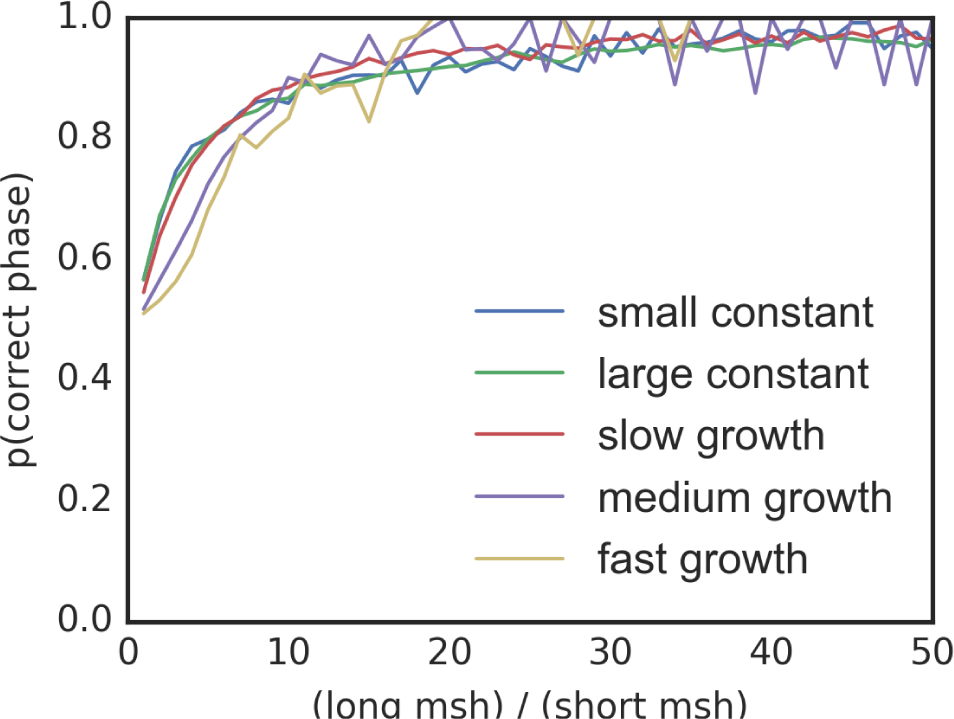
phasing accuracy of singleton variants as a function of the ratio of haplotype lengths under a variety of demographic models: samples of 1,000 chromosomes from constant population sizes of 10,000 and 1,000,000, and populations of 500,000 that have been growing exponentially at rates of 0.01%, 0.03%, and 0.1% per generation.

**Figure S4.**
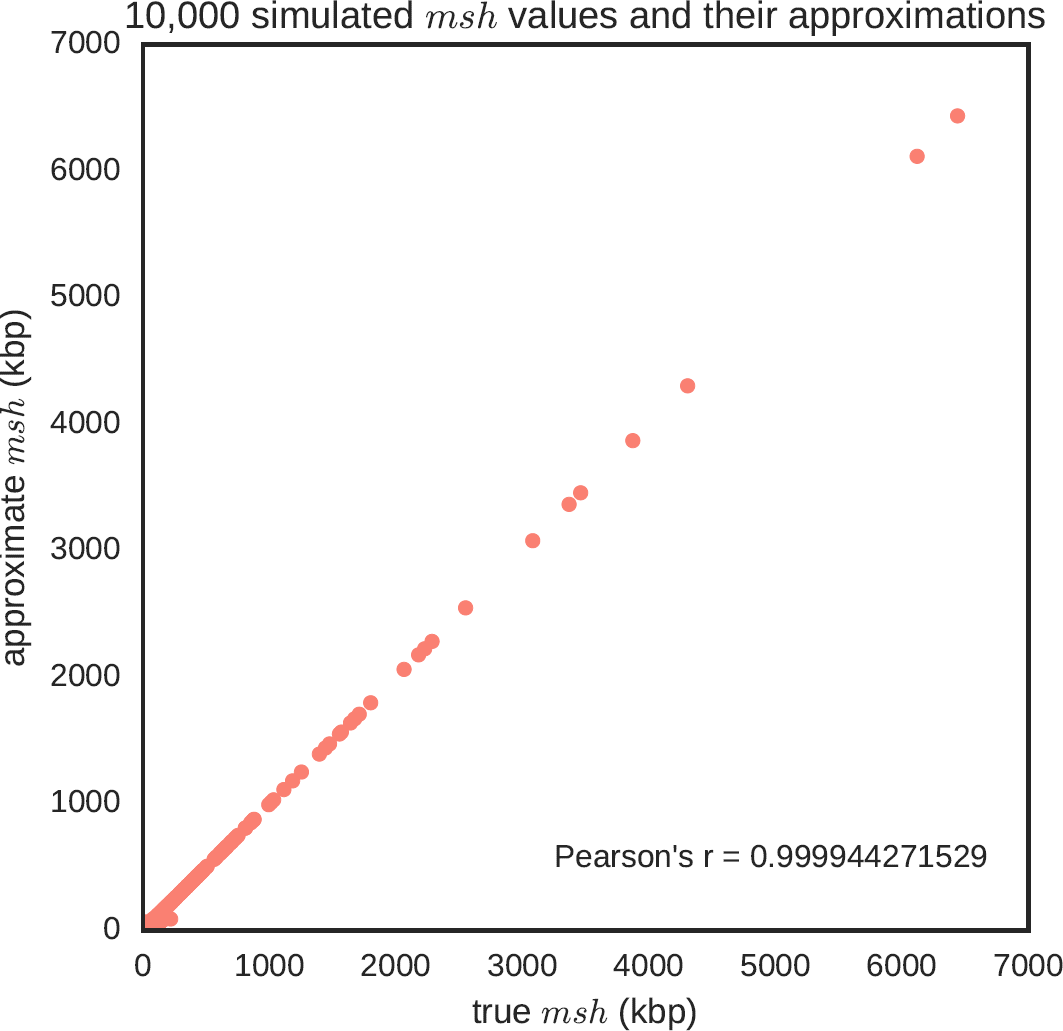
10,000 coalescent simulations comparing the *msh* of a singleton variant with the approximation of the *msh* considering only events on the external branch immediately ancestral to the singleton variant and its first sister branch. Each sample is 100 chromosomes drawn from a constant-sized diploid population of 1e6.

**Figure S5.**
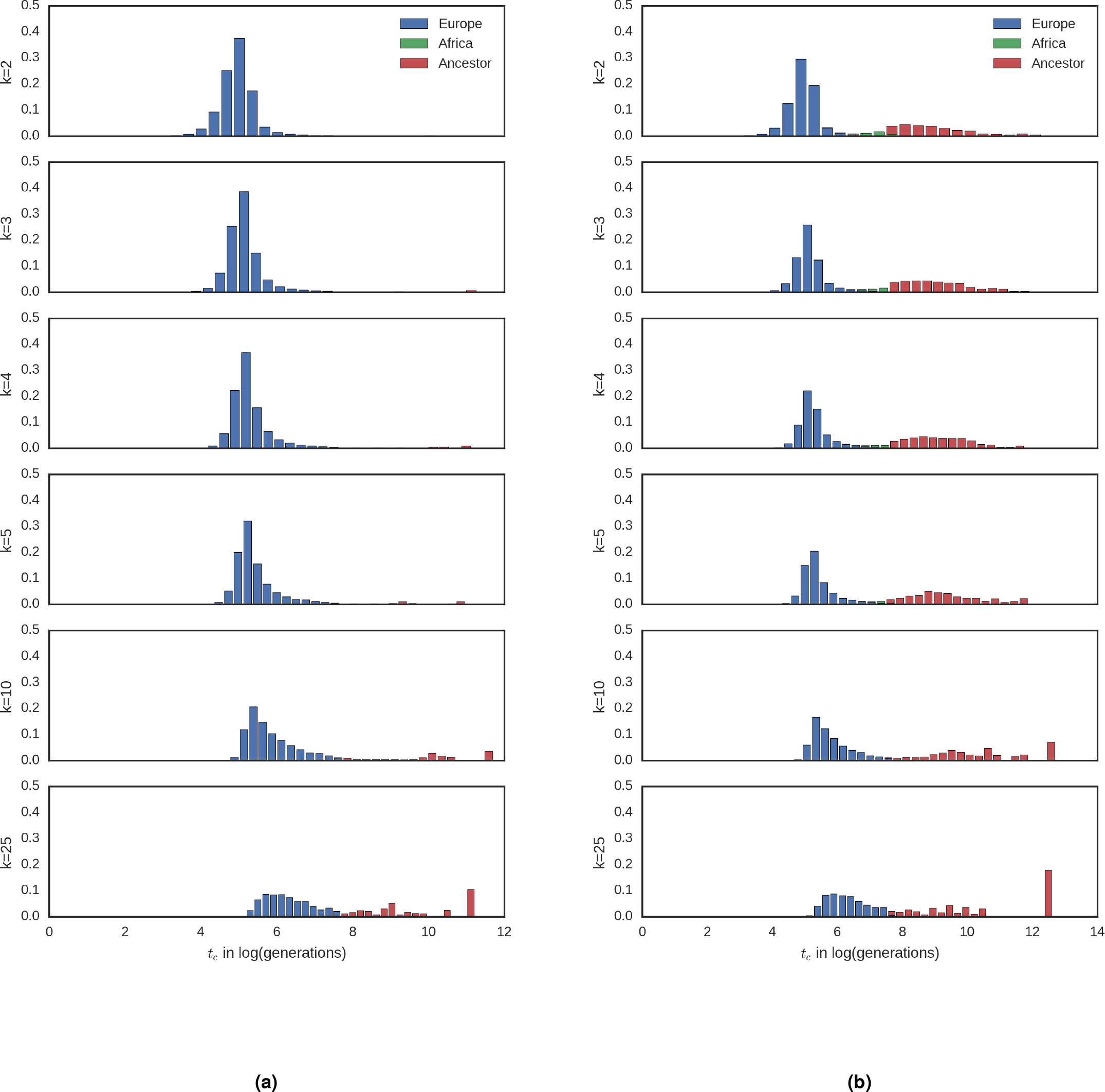
Expected distributions of *t_c_* values by age and population. Simulation results for the expected distribution of *t_c_* values colored by the population in which the coalescent event took place. Demographic models are (a) Gazave et al. and (b) African admixture (21gen).

**Figure S6.**
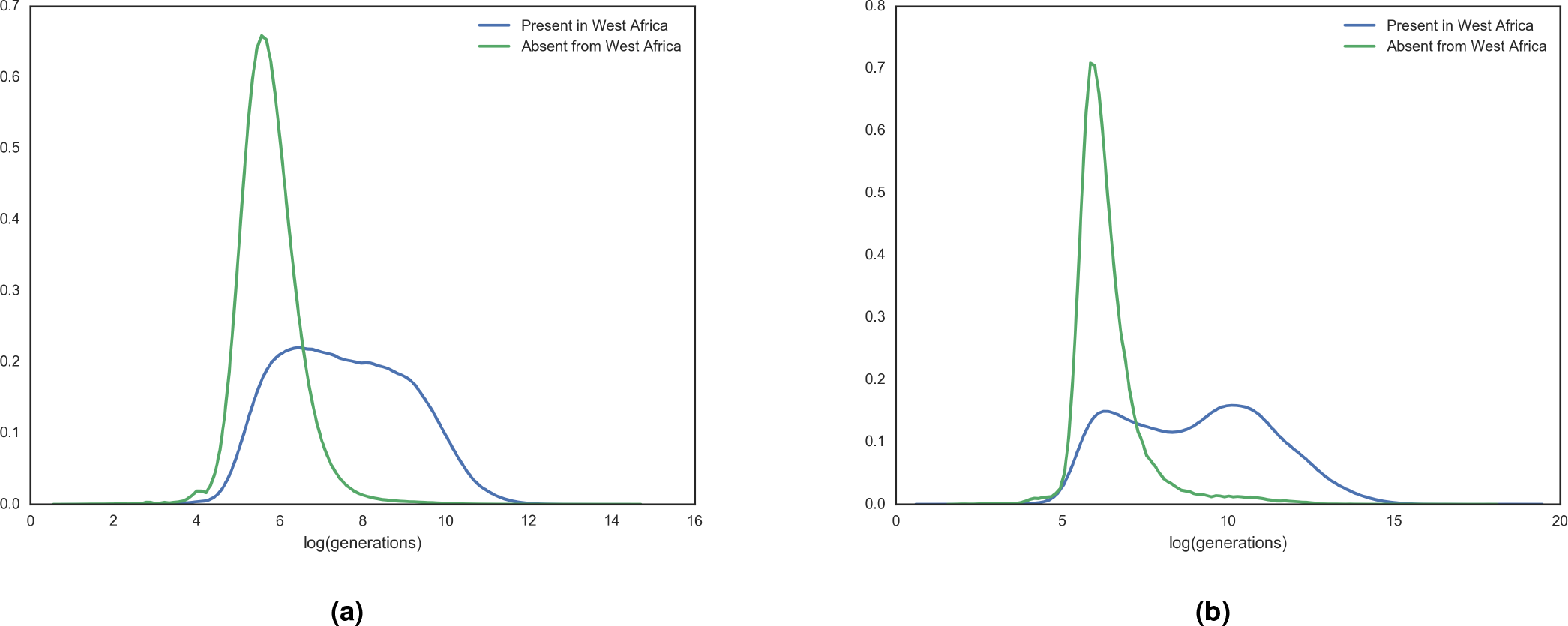
Age distribution of African variants. Distributions of *t_c_* values for UK10k singleton variants (a) and variants found 25 times (b) grouped by presence or absence in any West African population of the 1000 Genomes Project.

**Figure S7.**
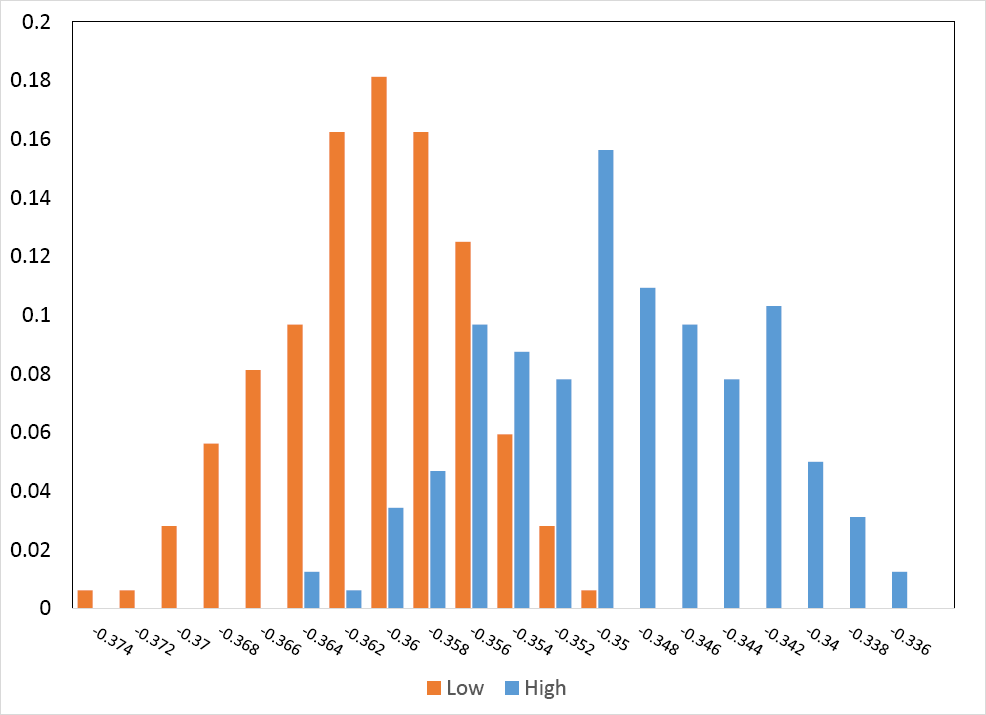
Histogram of *D* statistics calculated for UK10k individuals with the greatest (orange) and least (blue) number of singleton variants with 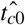 values *>* 3,000 generations.

**Figure S8.**
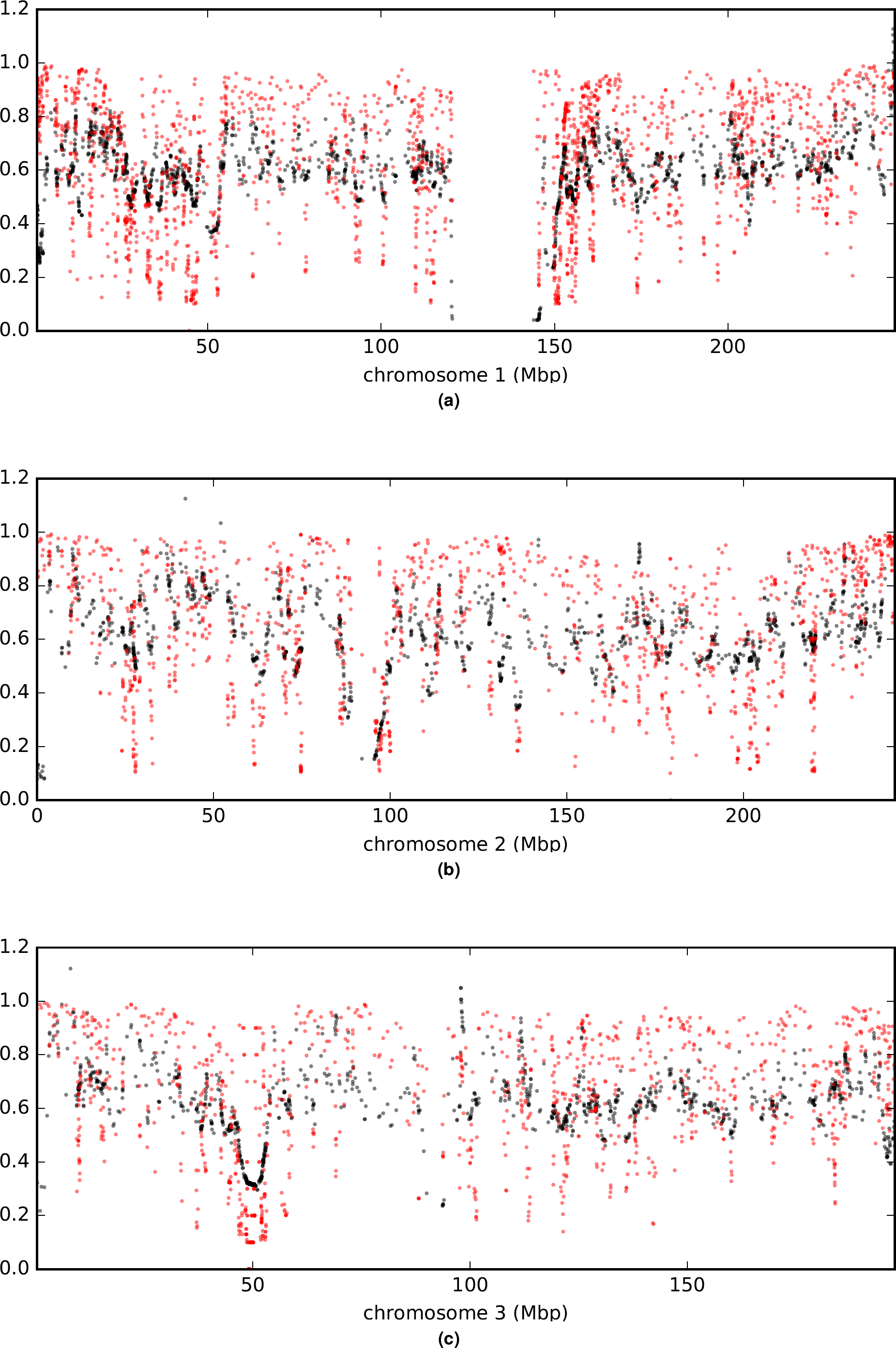
Comparison of *a* (rescaled 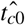; black) and *B* (in red) values across chromosomes.

**Figure S9.**
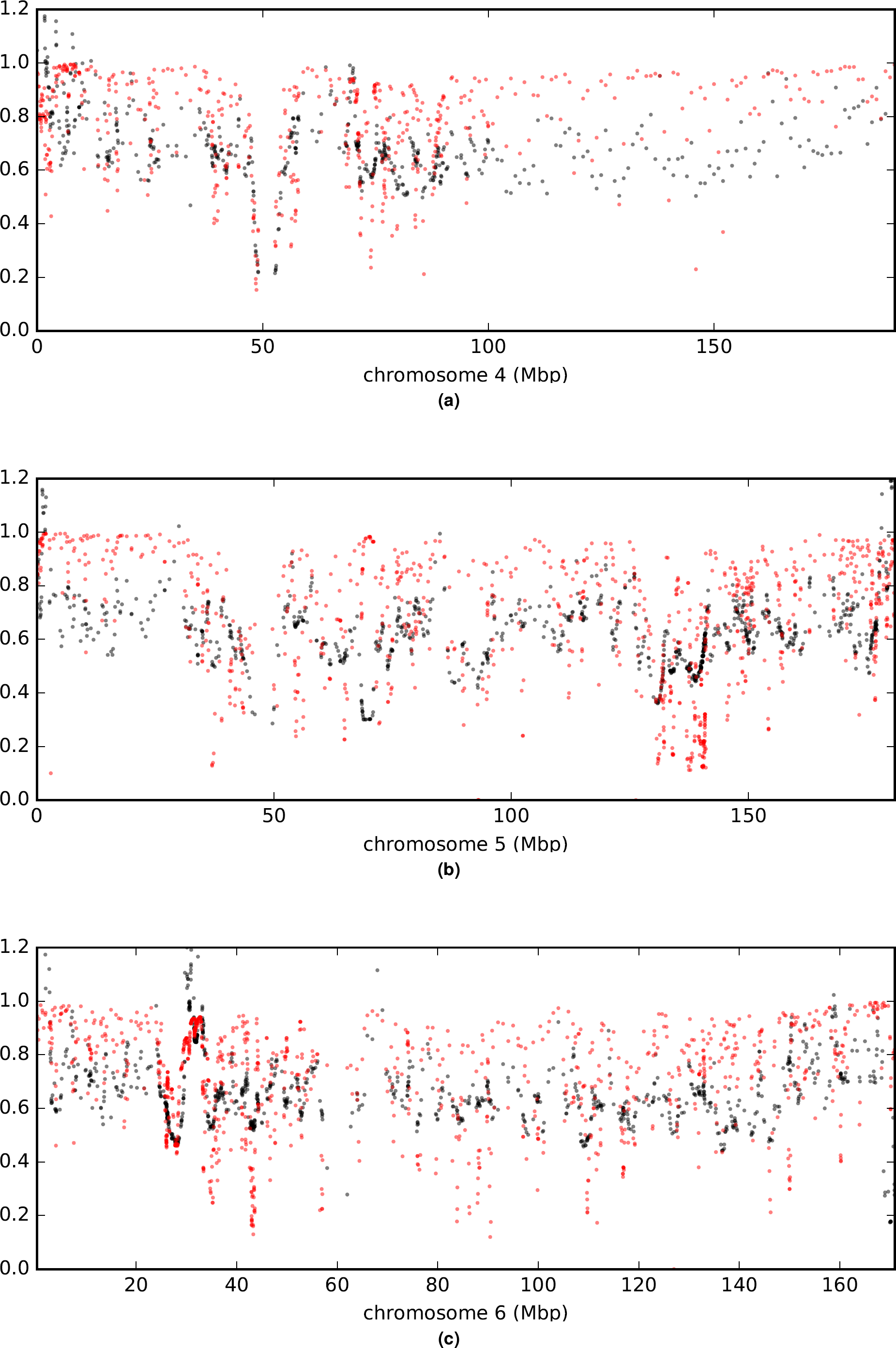
Comparison of *a* (rescaled 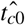; black) and *B* (in red) values across chromosomes.

**Figure S10.**
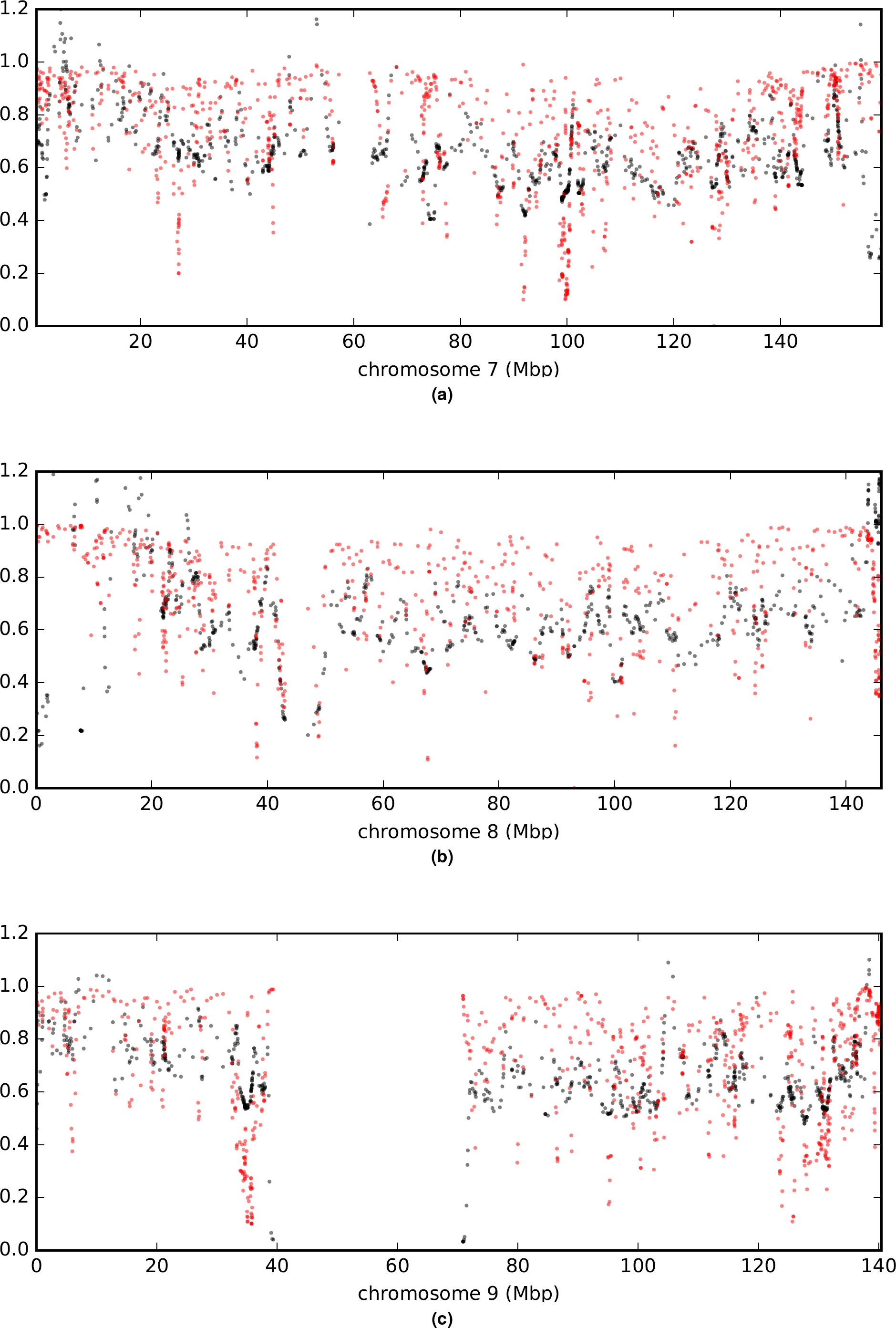
Comparison of *a* (rescaled 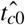; black) and *B* (in red) values across chromosomes.

**Figure S11.**
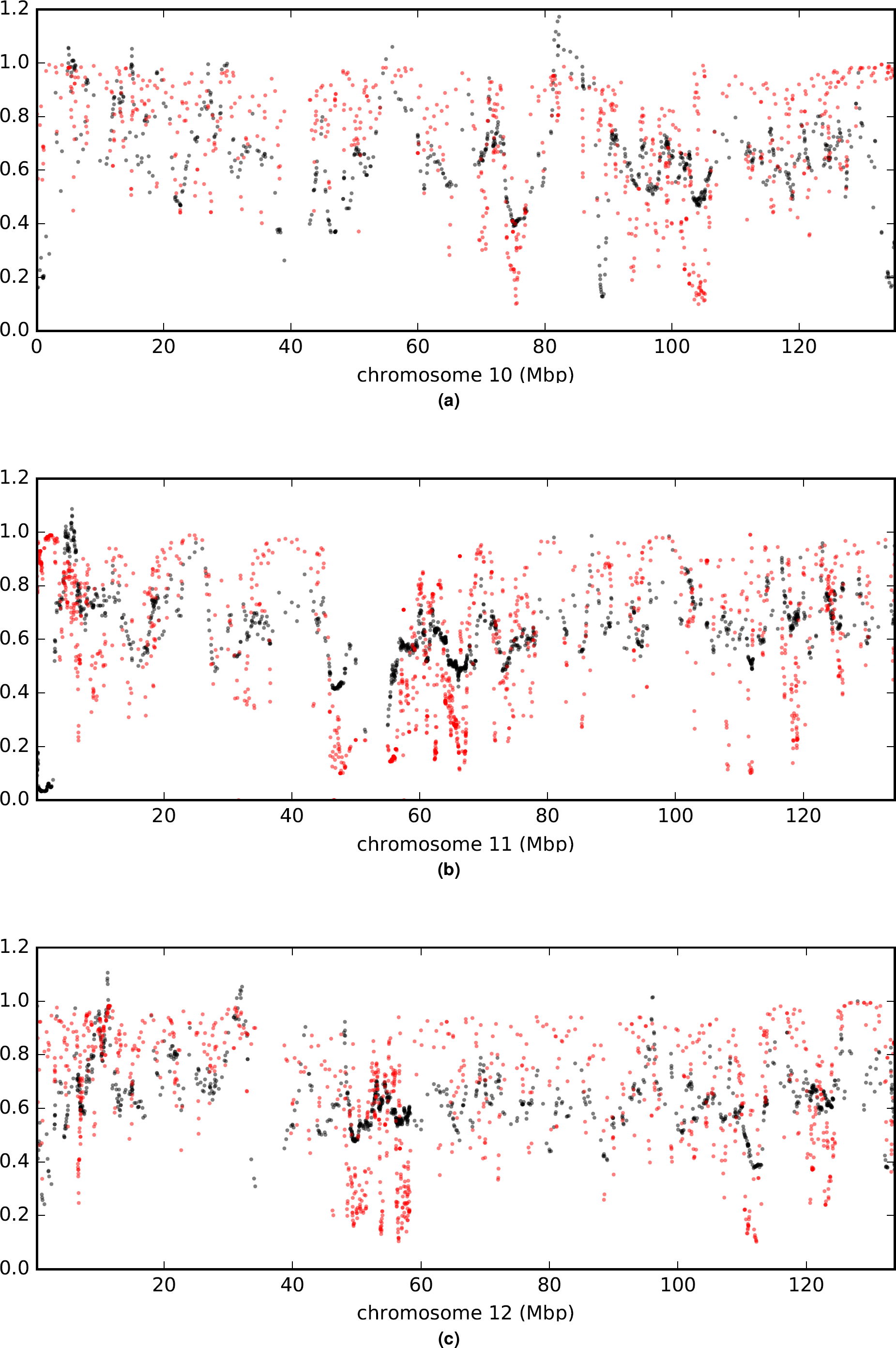
Comparison of *a* (rescaled 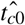; black) and *B* (in red) values across chromosomes.

**Figure S12.**
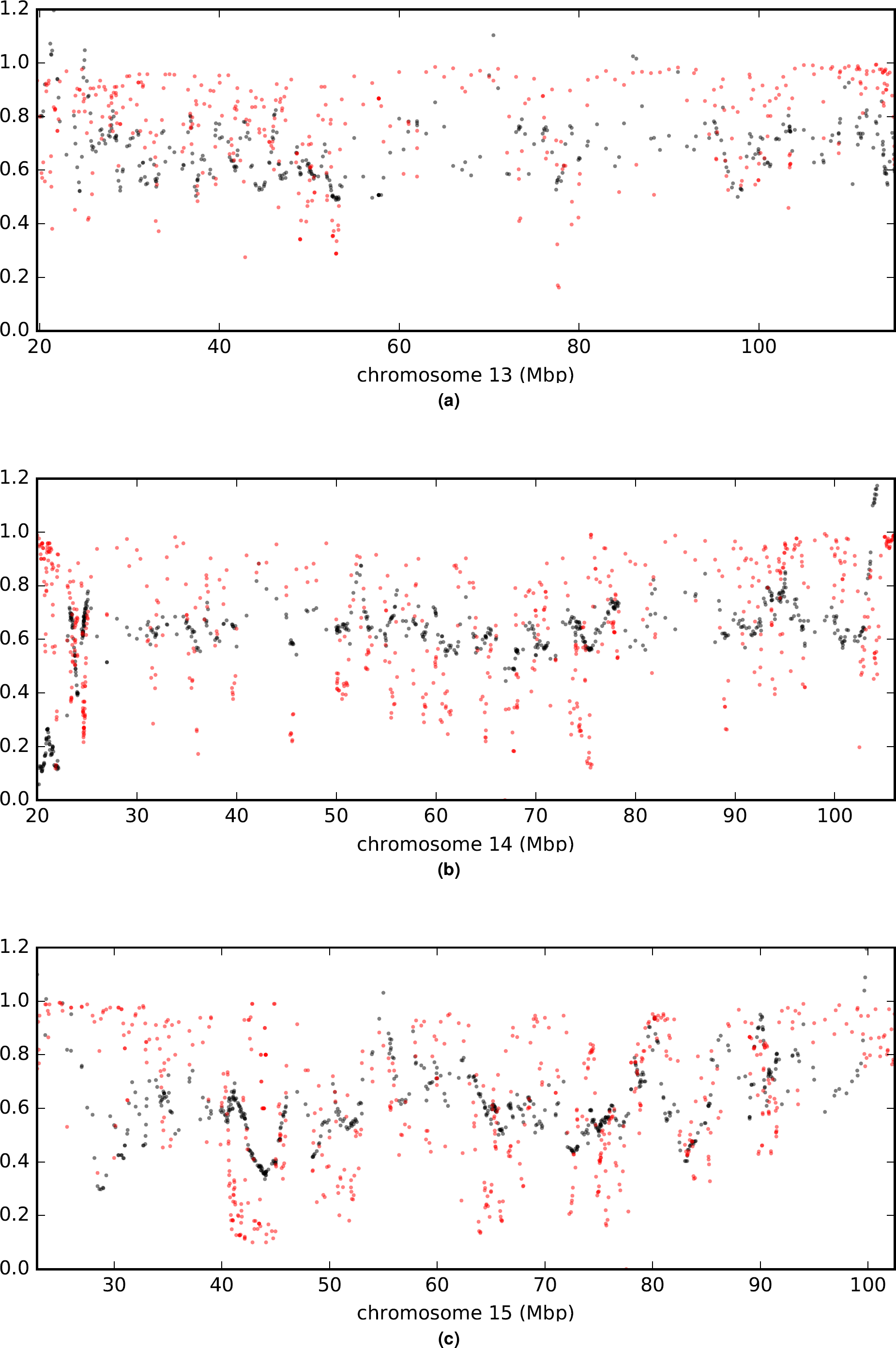
Comparison of *a* (rescaled 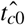; black) and *B* (in red) values across chromosomes.

**Figure S13.**
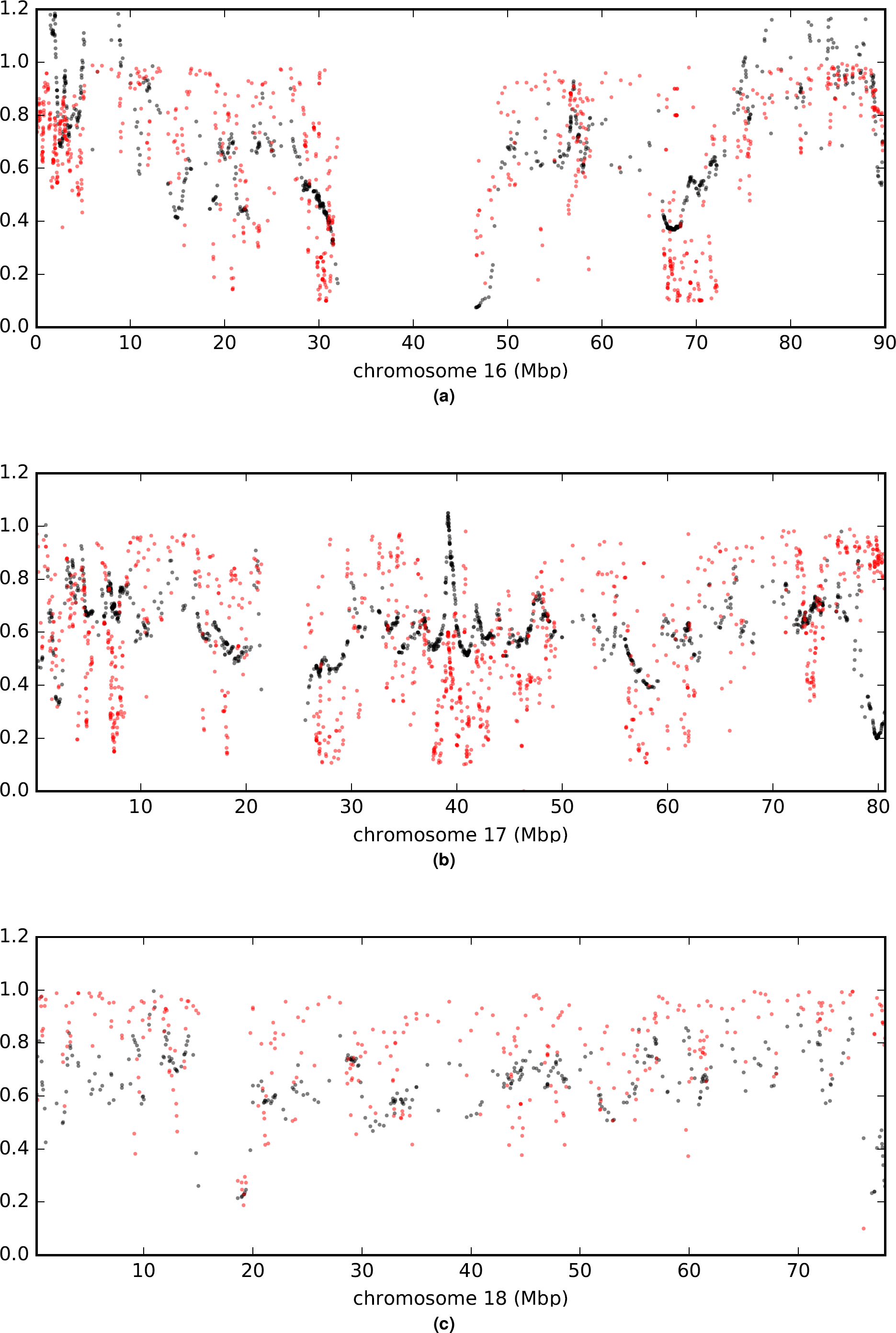
Comparison of *α* (rescaled 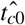; black) and *B* (in red) values across chromosomes.

**Figure S14.**
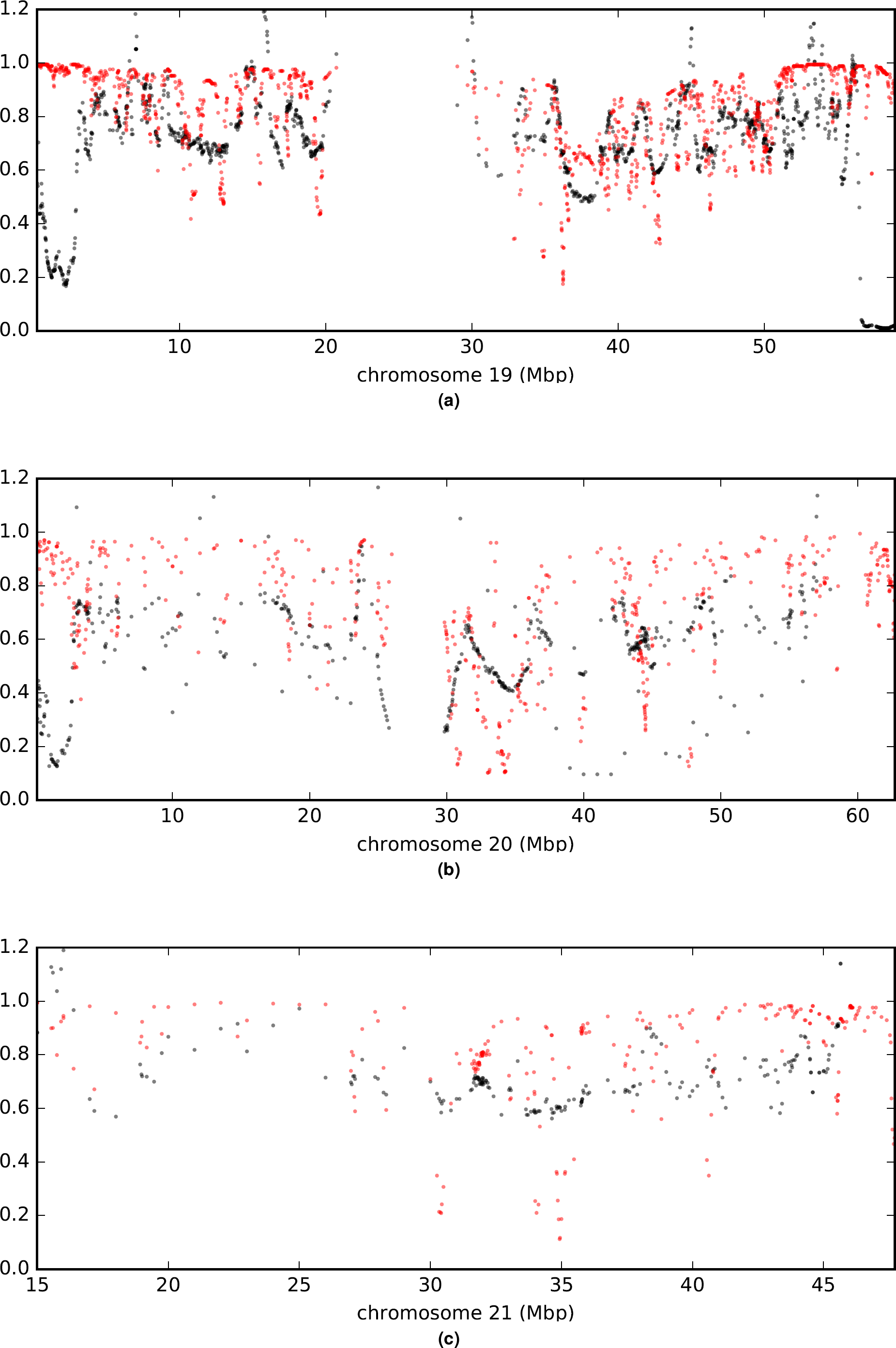
Comparison of *α* (rescaled 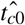; black) and *B* (in red) values across chromosomes.

## References

1. Maruyama, T. The age of an allele in a finite population. Genet. research 23, 137–143 (1974).

2. Reich, D. E. & Lander, E. S. On the allelic spectrum of human disease. TRENDS Genet. 17, 502–510 (2001).

3. Blekhman, R. et al. Natural selection on genes that underlie human disease susceptibility. Curr. biology 18, 883–889 (2008).

4. Luikart, G., Allendorf, F., Cornuet, J. & Sherwin, W. Distortion of allele frequency distributions provides a test for recent population bottlenecks. J. heredity 89, 238–247 (1998).

5. Andrés, A. M. et al. Targets of balancing selection in the human genome. Mol. biology evolution 26, 2755–2764 (2009).

6. Kidd, J. M. et al. Population genetic inference from personal genome data: impact of ancestry and admixture on human genomic variation. The Am. J. Hum. Genet. 91, 660–671 (2012).

7. Slatkin, M. & Rannala, B. Estimating the age of alleles by use of intraallelic variability. Am. journal human genetics 60, 447 (1997).

8. Mathieson, I. & McVean, G. Demography and the age of rare variants. PLoS Genet. 10, e1004528 (2014).

9. McVean, G. A. T. & Cardin, N. J. Approximating the coalescent with recombination. Philos. Transactions Royal Soc. B: Biol. Sci. 360, 1387–1393 (2005). URL http://rstb.royalsocietypublishing.org/content/360/1459/1387.abstract. DOI 10.1098/rstb.2005.1673.

10. Marjoram, P. & Wall, J. D. Fast” coalescent” simulation. BMC genetics 7, 16 (2006).

11. Paul, J. S., Steinrücken, M. & Song, Y. S. An accurate sequentially markov conditional sampling distribution for the coalescent with recombination. Genet. 187, 1115–1128 (2011).

12. Sabeti, P. C. et al. Detecting recent positive selection in the human genome from haplotype structure. Nat. 419, 832 (2002).

13. Field, Y. et al. Detection of human adaptation during the past 2000 years. Sci. aag0776 (2016).

14. The UK10K Consortium. The UK10k project identifies rare variants in health and disease. Nat. 526, 82–90 (2015). URL http://www.nature.com/nature/journal/v526/n7571/full/nature14962.html. DOI 10.1038/nature14962.

15. Gutenkunst, R. N., Hernandez, R. D., Williamson, S. H. & Bustamante, C. D. Inferring the joint demographic history of multiple populations from multidimensional SNP frequency data. PLoS Genet. 5, e1000695 (2009).

16. Gravel, S. et al. Demographic history and rare allele sharing among human populations. Proc. Natl. Acad. Sci. 108, 11983–11988 (2011). URL http://www.pnas.org/content/108/29/11983. DOI 10.1073/pnas.1019276108.

17. Gazave, E. et al. Neutral genomic regions refine models of recent rapid human population growth. Proc. Natl. Acad. Sci. 111, 757–762 (2014). URL http://www.pnas.org/content/111/2/757. DOI 10.1073/pnas.1310398110.

18. Gao, F. & Keinan, A. Inference of Super-exponential Human Population Growth via Efficient Computation of the Site Frequency Spectrum for Generalized Models. Genet. 202, 235–245 (2016). URL http://www.genetics.org/content/202/1/235. DOI 10.1534/genetics.115.180570.

19. Browning, S. R. & Browning, B. L. Accurate Non-parametric Estimation of Recent Effective Population Size from Segments of Identity by Descent. The Am. J. Hum. Genet. 97, 404–418 (2015). URL http://www.cell.com/ajhg/abstract/S0002-9297(15)00288-8. DOI 10.1016/j.ajhg.2015.07.012.

20. The 1000 Genomes Project Consortium. A global reference for human genetic variation. Nat. 526, 68–74 (2015). URL https://www.nature.com/nature/journal/v526/n7571/full/nature15393.html. DOI 10.1038/nature15393.

21. Green, R. E. et al. A draft sequence of the neandertal genome. Sci. 328, 710–722 (2010). URL http://www.sciencemag.org/cgi/content/abstract/328/5979/710. DOI 10.1126/science.1188021.

22. Patterson, N. et al. Ancient admixture in human history. Genet. 192, 1065–1093 (2012).

23. Kanehisa, M. & Goto, S. Kegg: kyoto encyclopedia of genes and genomes. Nucleic acids research 28, 27–30 (2000).

24. Tibshirani, R. Regression shrinkage and selection via the lasso. J. Royal Stat. Soc. Ser. B (Methodological) 267–288 (1996).

25. Platt, A., Weber, C. C. & Liberles, D. A. Protein evolution depends on multiple distinct population size parameters. BMC Evol. Biol. 18, 17 (2018). URL https://doi.org/10.1186/s12862-017-1085-x. DOI 10.1186/s12862-017-1085-x.

26. McVicker, G., Gordon, D., Davis, C. & Green, P. Widespread genomic signatures of natural selection in hominid evolution. PLoS genetics 5, e1000471 (2009).

27. Consortium, T. G. P. A map of human genome variation from population-scale sequencing. Nat. 467, 1061–1073 (2010). URL http://dx.doi.org/10.1038/nature09534http://www.nature.com/nature/journal/v467/n7319/abs/nature09534.html#supplementary-information.

28. Fenner, J. N. Cross-cultural estimation of the human generation interval for use in genetics-based population divergence studies. Am. J. Phys. Anthropol. 128, 415–423 (2005). DOI 10.1002/ajpa.20188.

29. Canny, N., Canny, N. P. & Low, A. *The Oxford History of the British Empire: Volume I: The Origins of Empire* (OUP Oxford, 2001). Google-Books-ID: eQHSivGzEEMC.

30. Shyllon, F. O. *Black People in Britain 1555-1833* (Institute of Race Relations, 1977). Google-Books-ID: IDVnAAAAMAAJ.

31. Kaufmann, M. *Africans in Britain: 1500-1640. Ph.D. thesis*, Thesis DPhil–University of Oxford (2011).

32. Moorjani, P. et al. The History of African Gene Flow into Southern Europeans, Levantines, and Jews. PLOS Genet. 7, e1001373 (2011). URL http://journals.plos.org/plosgenetics/article?id=10.1371/journal.pgen.1001373. DOI 10.1371/journal.pgen.1001373.

33. Botigué, L. R. et al. Gene flow from North Africa contributes to differential human genetic diversity in southern Europe. Proc. Natl. Acad. Sci. 110, 11791–11796 (2013). URL http://www.pnas.org/content/110/29/11791. DOI 10.1073/pnas.1306223110.

34. Nelson, M. R. et al. An abundance of rare functional variants in 202 drug target genes sequenced in 14,002 people. Sci. 337, 100–104 (2012).

35. Lancet, D. Exclusive receptors. Nat. 372, 321 (1994).

36. Alonso, S., López, S., Izagirre, N. & de la Rúa, C. Overdominance in the human genome and olfactory receptor activity. Mol. biology evolution 25, 997–1001 (2008).

37. Hoover, K. C. et al. Global survey of variation in a human olfactory receptor gene reveals signatures of non-neutral evolution. Chem. senses 40, 481–488 (2015).

38. O’Connell, J. et al. Haplotype estimation for biobank-scale data sets. Nat. genetics 48, 817 (2016).

39. Kelleher, J., Etheridge, A. M. & McVean, G. Efficient Coalescent Simulation and Genealogical Analysis for Large Sample Sizes. PLOS Comput. Biol. 12, e1004842 (2016). URL http://journals.plos.org/ploscompbiol/article?id=10.1371/journal.pcbi.1004842. DOI 10.1371/journal.pcbi.1004842.

40. Tennessen, J. A. et al. Evolution and Functional Impact of Rare Coding Variation from Deep Sequencing of Human Exomes. Sci. 337, 64–69 (2012). URL http://science.sciencemag.org/content/337/6090/64. DOI 10.1126/science.1219240.

41. Sharp, R. R. & Barrett, J. C. The environmental genome project: ethical, legal, and social implications. Environ. Heal. Perspectives 108, 279–281 (2000). URL http://www.ncbi.nlm.nih.gov/pmc/articles/PMC1638012/.

42. Cingolani, P. et al. A program for annotating and predicting the effects of single nucleotide polymorphisms, snpeff: Snps in the genome of drosophila melanogaster strain w1118; iso-2; iso-3. Fly 6, 80–92 (2012).

43. Hudson, R. R. & Kaplan, N. L. Deleterious background selection with recombination. Genet. 141, 1605–1617 (1995).

44. Nordborg, M., Charlesworth, B. & Charlesworth, D. The effect of recombination on background selection. Genet. Res. 67, 159–174 (1996).

